# Clinical extended-spectrum beta-lactamase antibiotic resistance plasmids have diverse transfer rates and can spread in the absence of antibiotic selection

**DOI:** 10.1101/796243

**Authors:** Fabienne Benz, Jana S. Huisman, Erik Bakkeren, Joana A. Herter, Tanja Stadler, Martin Ackermann, Médéric Diard, Adrian Egli, Alex R. Hall, Wolf-Dietrich Hardt, Sebastian Bonhoeffer

## Abstract

Horizontal gene transfer, mediated by conjugative plasmids, is a major driver of the global spread of antibiotic resistance. However, the relative contributions of factors that underlie the spread of clinically relevant plasmids are unclear. Here, we quantified conjugative transfer dynamics of Extended Spectrum Beta-Lactamase (ESBL) producing plasmids in the absence of antibiotics. We showed that clinical *Escherichia coli* strains natively associated with ESBL-plasmids conjugate efficiently with three distinct *E. coli* strains and one *Salmonella* enterica serovar Typhimurium strain, reaching final transconjugant frequencies of up to 1% within 24 hours *in vitro*. The variation of final transconjugant frequencies varied among plasmids, donors and recipients and was better explained by variation in conjugative transfer efficiency than by variable clonal expansion. We identified plasmid-specific genetic factors, specifically the presence/absence of transfer genes, that influenced final transconjugant frequencies. Finally, we investigated plasmid spread within the mouse intestine, demonstrating qualitative agreement between plasmid spread *in vitro* and *in vivo.* This suggests a potential for the prediction of plasmid spread in the gut of animals and humans, based on *in vitro* testing. Altogether, this may allow the identification of resistance plasmids with high spreading potential and help to devise appropriate measures to restrict their spread.

## Introduction

Plasmids can transfer horizontally between bacterial cells of the same or of different species, thereby leading to rapid spread of accessory genes and allowing bacterial populations to adapt to new environments. Antibiotic resistance determinants are often plasmid-encoded and resistance plasmids drive the continuous rise of resistant bacterial pathogens that undermine the effectiveness of antibiotics in the clinic (1–3). Whether a plasmid can increase in frequency within a bacterial population depends on the plasmid as well as its bacterial host. Some plasmids become global threats in association with a specific clone (4–6). Other plasmids, however, are a clinical problem independent of their host strain (7). The family of Enterobacteriaceae is a key public health concern, as it includes some of the most important nosocomial pathogens and pandemic extended-spectrum beta-lactamases (ESBLs) producing strains (4,8–10). ESBLs provide their bacterial host with resistance to beta-lactam antibiotics, such as the widely used penicillins and cephalosporins, and are mostly encoded on plasmids of the incompatibility groups IncF and IncI (9,11,12).

*In vitro* studies revealed several key factors driving changes in plasmid frequency over time. For instance, if a plasmid of the same incompatibility group is already established in a potential recipient, plasmid incompatibility and surface- or entry exclusion may inhibit further plasmid acquisition (13, 14). Some co-residing plasmids can enhance each other’s stability (15) and allow or increase their transfer to a new bacterial host (16, 17). Once a plasmid has been taken up by a bacterial cell, its replication system and the interaction with other host factors define whether it can be replicated and stably maintained. Bacterial immunity systems such as CRISPR, or a restriction modification system (RM system) different from the ones in the previous bacterial host, can eliminate an incoming plasmid (18–21). Whether a plasmid persists in a bacterial population over a longer time period depends on the frequency of plasmid loss during host replication (22) and the cost of plasmid carriage (23), although the latter can be reduced by the accumulation of compensatory mutations allowing for plasmid persistence (24). Such persistence can also occur when the cost of a plasmid in terms of a growth disadvantage is outperformed by its high rates of conjugation (25).

Save for a notable exception (26), these factors have mostly been studied separately from each other, with plasmids not directly relevant for antibiotic resistance spread or with individual plasmids that have been moved into well-defined model strains (18,25,27). But to better understand the drivers of the spread of antibiotic resistance plasmids, we need to learn more about these factors’ relative contributions to plasmid spread among clinical strains and in relevant environments. The gut of humans and farm animals contain a dense microbiota, which represent hotspots for bacterial interactions and transfer of antibiotic resistance plasmids. This knowledge relies on genomic studies (28–30) which, however, do not allow an investigation of the dynamics and drivers of plasmid spread. We, and others (31), have previously used mouse models to study processes that limit or boost plasmid spread in the gut (32–36). These studies, however, were limited to laboratory strains and single conjugative plasmids without clinical relevance, with the exception of one ESBL-plasmid used in (34). To our knowledge, there are no studies that compare plasmids and their transfer dynamics quantitatively *in vivo*.

Here, we use clinical *E. coli* strains and their natively associated ESBL-plasmids. We combine bioinformatic analyses with *in vitro* and *in vivo* experiments to provide a comprehensive and quantitative investigation of plasmid spread in the absence of antibiotic selection. Our *in vitro* conjugation experiments showed that the final frequencies of ESBL-plasmid carrying recipient strains (transconjugants) varied largely and were determined by plasmid, donor and recipient factors. The plasmid itself had the largest effect: Of the plasmids we tested, only those with functional transfer (*tra*) genes, including those encoding the sex pilus and the proteins required for the conjugative transfer (37–39), transferred to multiple recipients. None of the ESBL-plasmids lead to significant growth costs for recipient strains, neither *in vitro* nor *in vivo*. The plasmid frequency in recipient populations was determined by the rate at which they transferred horizontally to recipient strains (transfer rate). The rates of plasmid transfer varied depending on the donor-plasmid pair and the recipient strain. Importantly, our *in vitro* testing qualitatively predicted the plasmid spread in our antibiotic-free murine model for gut colonization. Furthermore, plasmid carriage was not associated with a cost in the gut and horizontal plasmid transfer determined the frequency of transconjugants. With this study, we contribute to connecting mechanisms of plasmid spread found *in vitro* with corresponding patterns *in vivo* and ultimately to a better understanding of the factors underlying the effective spread of clinically relevant resistance plasmids (5, 40).

## Materials and methods

### Strains and growth conditions

We used 8 ESBL-plasmid positive *E. coli* strains as plasmid donors (D1-D8; Supplementary Table 1). They were sampled from patients in a transmission study at the University Hospital Basel, Switzerland, and their ESBL-plasmids reflect relevant vectors of ESBL mediated drug resistance (41). This collection comprised strains belonging to sequence types (ST) ST117, ST648, ST40, ST69, ST80, ST95, ST6697 and the very common ESBL sequence type ST131. We worked with 4 ESBL-plasmid negative recipient strains: RE1, a mouse-derived *E.coli* strain cured of its native IncI1 plasmid (35); RE2 and RE3, two clinical *E. coli* isolates from healthy patients (41, 42); and RS1, the *Salmonella enterica* Typhimurium strain ATCC 14028 (RS). A comprehensive list of the plasmids found in these strains is given in Supplementary Table S1. Marker plasmids were introduced by electroporation, to mark recipients with either pACYC184 (New England Biolabs) encoding Chloramphenicol (Cm) resistance (except for RS, having chromosomal Cm resistance *marT::cat* (32)) or pBGS18 (43) encoding Kanamycin (Kan) resistance. Unless stated otherwise, we grew bacterial cultures at 37°C and under agitation (180 rpm) in lysogenic broth (LB) medium, supplemented with appropriate amounts of antibiotics (none, 100µg/mL Ampicillin (Amp), 25 µg/mL Cm *in vitro* and 15µg/mL Cm prior to *in vivo* experiments, 50µg/mL Kan). We stored isolates in 25% glycerol at −80°C.

### Antibiotic resistance profiling

We used microdilution assays with a VITEK2 system (bioMérieux, France) to determine the minimum inhibitory concentrations (MIC). MIC breakpoints for ESBLs were interpreted according to EUCAST guidelines (v8.1). In addition, we confirmed resistance mechanism phenotypically, using ROSCO disk assays (ROSCO Diagnostica, Denmark), and/or genotypically with detection of CTX-M1 and CTX-M9 groups using the eazyplex Superbug assay (Amplex, Germany).

### *In vitro* conjugation experiment

We determined plasmid spread as the final frequency of the recipient population that obtained an ESBL-plasmid (transconjugants/(recipients+ transconjugants), T/(R+T)) in a high throughput, 96-well plate-based assay. Donor and recipient populations grew over night with or without Amp, respectively. We washed the independent overnight cultures by spinning down and resuspending and added ∼1µL of 6.5-fold diluted donor and recipient cultures into 150µL fresh LB with a pin replicator (total ∼1000-fold dilution, aiming to reach approximately a 1:1 ratio of donor and recipient). These mating populations grew for 24 hours in the absence of antibiotics and were only shaken prior to hourly optical density (OD) measurements (Tecan NanoQuant Infinite M200 Pro). To determine the final cell densities, we plated the mating cultures at the end of the conjugation assay on selective LB-plates. In the first conjugation experiment, referred to as the 1^st^ generation *in vitro* experiment, where the clinical strains transferred their native plasmids to recipients, we selected for donors+transconjugants with Amp, for recipients+transconjugants with Cm (*E. coli* recipients carried pACYC184-Cm and RS chromosomal *marT::cat*) and for transconjugants with Amp+Cm. For a second conjugation experiment, referred to as the 2^nd^ generation *in vitro* experiment, we chose a subset of transconjugants generated in the 1^st^ generation *in vitro* experiment as new plasmid donors. Transconjugants and recipients of the clone type RE3 were omitted because of the size of the experiment, and transconjugant RE2 carrying p1B_IncI had to be excluded as plasmid donor due to insufficient freezer stocks. We selected for donors with Cm, for recipients with Kan (recipients carried pBGS18-Kan) and for transconjugants with Kan+Amp. With this approach, we were able to detect transconjugant populations if their fraction exceeded 10^-8^ cells /mL. We performed experiments with *E. coli* recipients and *S.* Typhimurium recipient RS as independent experiments and the 1^st^ generation *in vitro* (*n=4-6*) and 2^nd^ generation *in vitro* (*n=6*) experiments each in two replica blocks.

The plasmids in our conjugation experiments could potentially be horizontally transferred either in the liquid mating population or after plating on selective plates (surface-mating). To assess the extent of surface-mating we performed an additional experiment, where we treated donors and recipients as above but grew them separate liquid cultures, instead of mixed cultures, only mixing them immediately before plating on selective LB-plates. To compare plasmid spread of 1^st^ generation and 2^nd^ generation *in vitro* experiments side by side, we performed a conjugation experiment as described above with D1 (with and without pACYC184) and transconjugants RE1 and RS carrying plasmid 1B_IncI, isolated from the 1^st^ generation *in vitro* experiment, as plasmid donors.

### *In vitro* plasmid cost experiment and other growth rate measurements

To investigate the effect of ESBL-plasmid carriage on bacterial growth in absence of antibiotics, we measured the growth rate of transconjugants and recipients. Per donor and recipient combination, we used three transconjugants, four replicates each, obtained from independent mating populations of the 1^st^ generation *in vitro* experiment. Transconjugants for which we have not stored three independent transconjugants were excluded from this analysis. We grew bacterial cultures in absence of antibiotics overnight and diluted them 150-fold by transfer with a pin replicator to a 96-well plate, containing 150µL fresh LB per well. We incubated the cultures without shaking and estimated growth rates of recipients and transconjugants based on ten manual OD measurements over 24 hours. We estimated growth rates (h^-1^) using the R package Growthcurver (44). We expressed plasmid cost as the growth rate of transconjugants relative to the corresponding ESBL-plasmid free recipient. Transconjugants have experienced longer growth under laboratory conditions than recipients (conjugation experiment). To verify that this did not affect our estimates of plasmid cost, we conducted a third growth rate experiment. First, recipients were grown under the same conditions as in the conjugation experiment but in absence of donor strains. Second, the growth rate of these treated strains was compared to the growth rates of untreated recipients (Supplementary Figure S1).

Other growth rate measurements (Supplementary Figures S1, S16) were performed as follows: We grew bacterial cultures with appropriate antibiotics overnight, washed and diluted them ∼1000-fold. For the growth measurements, bacteria grew in the absence of antibiotics and the plate reader measured OD every hour for 24 hours. Again we estimated growth rates (h^-1^) using the R package Growthcurver (44).

### *In vivo* experiments

We have previously established a murine model for enterobacterial pathogen infection (45) that allows to detect plasmid spread (32–36). For conjugation experiments, we used 8-16 week old C57BL/6 mice that contain an oligo microbiota allowing colonization of approximately 10^8^ *E. coli* per gram faeces (46). *E. coli* stool densities of up to 10^8^ cfu/g have also been detected in healthy human volunteers (42). We infected 7-10 mice per treatment group (minimum of two independent experiments; no antibiotic pre-treatment) orogastrically with ∼5×10^7^ CFU of RE2 or RE3, carrying marker plasmid pACYC184 and 24 hours later with ∼5×10^7^ CFU of either D4, D7, or D8. Faeces were collected daily, homogenized in 1 ml of PBS with a steel ball by a Tissue Lyser (Qiagen) at 25 Hz for 1 min. We enumerated bacterial populations by selective plating on MacConkey media (selection for donors+transconjugants with Amp (100 µg/mL), for recipients+transconjugants with Cm (15 µg/mL) and for transconjugants with Amp+Cm) and calculated final transconjugant frequencies T/(R+T).

For competition experiments we infected 8-16-week-old C57BL/6 oligo microbiota mice orogastrically with a 1:1 mixture of both competitor strains (∼5×10^7^ CFU total; no antibiotic pre-treatment). We collected faeces and enumerated bacterial populations daily. We plated bacteria on MacConkey agar containing Cm and replica-plated on media containing Cm, Kan, and Amp to select the transfer deficient transconjugants. A change in fitness conferred by plasmid carriage is reflected in the relative frequency of recipients to transconjugants (R/T).

Prior to all infections, we subcultured the overnight cultures (LB containing the appropriate antibiotics) for 4 hours at 37°C without antibiotics (1:20 dilution) to ensure equal densities of bacteria. Cells were washed in PBS and introduced into mice. All infection experiments were approved by the responsible authority (Tierversuchskommission, Kantonales Veterinäramt Zürich, license 193/2016 and license 158/2019).

### Sequencing, assembly, annotation

We sequenced all donor and recipient strains with Illumina MiSeq (paired end, 2×250 bp), Oxford Nanopore MinION and PacBio Sequel methods. We produced hybrid assemblies with Unicycler (47) (v0.4.7) and used the most contiguous assemblies (Oxford Nanopore – Illumina for D1,D2,D4,D6,D7,D8 and Pacbio Sequel – Illumina for D3,D5, RE1, RE2, RE3). Manual curation involved removing contigs smaller than 1kB, and sequences up to 5 kB that mapped to the own chromosome. We performed quality control by mapping the paired end Illumina reads to the finished assemblies using samtools (v1.2) and bcftools (v1.7) (48, 49). For recipient RS, the ancestral strain was sequenced with Illumina (2×150 bp), and mapped against the reference sequence, downloaded from NCBI Genbank under the accession numbers NZ_CP034230.1 and NZ_CP034231.1.

To study the genetic contribution to the observed variation in plasmid spread, we sequenced various transconjugants from the 1^st^ and 2^nd^ generation *in vitro* experiments as well as the *in vivo* transfer experiment (Supplementary Table S2). *In vitro*: three clones from independent mating populations for RE3 carrying plasmid p4A_IncI or p8A_IncF and one clone for the other transconjugants. *In vivo*: eight clones of RE3 carrying p4A_IncI isolated from five mice on day 7 post donor infection, and eight clones of RE3 carrying p8A_IncF isolated from six mice on day 2 (1 clone), day 6 (3 clones), or day 7 (4 clones) post donor infection. Resequencing was performed on an Illumina MiSeq (paired end, 2×150 bp) and we mapped the reads to the closed assemblies of respective donor and recipient strains using the breseq pipeline (v 0.32.0) (50). Mutations or indels shared by all re-sequenced strains were treated as ancestral (Supplementary Table S2).

To investigate the transfer of plasmid p8C_IncBOKZ, we screened 3-5 transconjugants from independent mating populations per conjugation pair (Supplementary Table S2). We performed PCRs with primers specific to IncB/O and IncK plasmids (51), (5′ to 3′): MRxeBO_K_for: GAATGCCATTATTCCGCACAA and MRxeBO_K _rev; GTGATATACAGACCAT-CACTGG).

To extract a chromosomal alignment of the *E. coli* donor and recipient strains, we concatenated the genes returned by core genome Multi-Locus Sequence Typing (cgMLST) for all strains. We used the chewBBACA software to type all strains according to the Enterobase cgMLST scheme (52, 53). We inferred the phylogenetic tree using BEAST2 (54), with an HKY substitution model, a fixed mutation rate (fixed to the *E. coli* mutation rate of 10^-4^ mutations per genome per generation, as estimated by Wielgoss et al. (55)), and a birth-death tree prior (priors are listed in Supplementary Table S4). We performed bacterial genome annotation using Prokka (56), and determined the sequence type (ST) using mlst (Torsten Seemann, https://github.com/tseemann/mlst), which makes use of the PubMLST website (https://pubmlst.org/) developed by Keith Jolley (57). Phylogroups were assigned using ClermonTyper (58).

We determined genomic features using a range of bioinformatic tools, and by BLAST comparison against various curated databases. Plasmid replicons and resistance genes were identified using abricate (Torsten Seemann, https://github.com/tseemann/abricate) with the PlasmidFinder (59) and ResFinder (60) databases respectively. We located phages using PHASTER (61) (listing only those marked as “complete”), type 6 secretion systems using SecReT6 (62), virulence genes using the Virulence finder database (63), toxin-antitoxin systems using the database TADB 2.0 (64), and CRISPR-Cas loci using CRISPRCasFinder (65). We found restriction-modification (RM) systems using grep on the term ‘restriction’ in the general feature format (GFF) files from prokka, and verified them with the RM-database Rebase (66). To determine the presence/absence of IncF and IncI transfer genes, we constructed our own database as a reference. IncF transfer genes were taken from the supplementary material of Fernandez-Lopez et al. (67), IncI1 transfer genes from plasmids R64 using the annotations by Komano et al. (37), and Inc1γ transfer genes from the plasmid R621a annotated by Takahashi et al. (38).

### Construction of non-transferrable plasmids

For the *in vivo* competition experiments, we generated non transferrable plasmids for three independent transconjugants. We deleted their origin of transfer (*oriT*) region using the lambda red recombinase system with pKD4 as template for the Kan resistance marker (68). The following primers were used (5′ to 3′): For IncI plasmids (p4A_IncI) DIncI_oriTnikA_f (GCATAAGACTATGATGCACAAAAATAAC-AGGCTATAATGGGTGTAGGCTGGAGCTGCTTC) and DIncI_oriTnikA_r (CCTTCTCTTTTTCG-GAATGACTGCATTCACCGGAGAATCCATGGGAATTAGCCATGGTCC) (35) and for F plasmids (p8A_IncF) D25_2_oriT-nikA-ko_vw (CCATGATATCGCTCTCAGTAAATCCGGGTCTATTTTGTA-AGTGTAGGCTGGAGCTGCTTC) and D25_2_oriT-nikA-ko-rev (GTGCGGACACAGACTGGATATTT-TGCGGATAAAATAATTTATGGG-AATTAGCCATGGTCC). We verified all mutants by PCR (IncI1_oriT _val_f: AGTTCCTCA-TCGGTCATGTC, IncI1_oriT _val_r: GAAGCCATTGGCACTTTCTC, D25_oriT _val_fw: CATACAGG-GATCTGTTGTC and D25_2_oriT_ver_rv: CAGAATCACTAT-TCTGACAC) and experimentally by loss of transfer function.

### Statistical analyses

For *in vitro* experiments we performed analyses using R (version 3.4.2). The effects of donor, recipient and plasmid on final transconjugant frequency were analyzed with either a two-way ANOVA (1^st^ generation *in vitro* experiment with factors donor-plasmid pair and recipient) or a three-way ANOVA (2^nd^ generation *in vitro* experiment with factors donor, plasmid, recipient). For the 1^st^ generation *in vitro* experiment, we excluded strain-plasmid pairs which did not result in transconjugants (D2, D3, D7) and recipient RS from this analysis. When single replicates for a given donor-recipient combination lacked transconjugants (D5 and D6), we assigned these replicates a final transconjugant frequency at the detection limit of 10^-8^. With the same data we performed the correlation of final transconjugant frequency and transfer rate. The data of the 2^nd^ generation *in vitro* experiment was not fully factorial. To enable testing of interactions, we therefore performed two 3-way ANOVAs: one excluding plasmid p1B_IncI and one excluding donor RE2, for which we had to take the two replicate blocks into account: *P* < 0.001). For two replicate populations (RS self-self transfer with p1B_IncI), we had higher counts on plates selecting for transconjugants than on plates selecting for recipients+transconjugants and replaced the resulting negative CFU/mL for recipients with 0 CFU/mL (we assume the higher count on selective agar reflects measurement error, given the true frequency of plasmid-carrying cells cannot exceed 1.0). We used the end-point method of Simonsen *et al.* to determine plasmid transfer rates in diverse bacterial populations (69).

For statistical comparisons derived from *in vivo* experiments, Kruskal-Wallis tests were performed with Dunn’s multiple test correction using GraphPad Prism Version 8 for Windows.

## Results

### Strains and plasmids

We used eight clinical E. coli strains as ESBL-plasmid donors (D1-D8) and four recipient strains susceptible to β-Lactam antibiotics, of which three are E. coli (RE1-RE3) and one S. Typhimurium (RS, Supplementary Figure S2, Supplementary Table S3). Sequence analysis revealed a large phylogenetic diversity, with donor strains belonging to phylogenetic groups B1, B2, D or its subgroup F, and recipients to either B2 or A (Figure 1). We also observed diverse accessory traits such as bacterial immunity systems (Supplementary Figure S3) and virulence genes (Supplementary Figure S4). All but two strains encode type 6 secretion systems (T6SS), and strain D4 shows an Enteropathogenic (EPEC) virulence profile. Each strain carries at least one and up to eight plasmids of various incompatibility groups (Supplementary Figure S5, Table S1). Every donor strain harbours a single antibiotic-resistance plasmid (the ESBL plasmid), either of the plasmid family IncI or IncF (Table 1) and displayed an ESBL-resistance phenotype (Supplementary Table S3). All strains encode numerous intact prophage sequences in their chromosome (Supplementary Figure S6) and we found P1-like phages, i.e. prophages that move like plasmids in their lysogenic phase (70, 71), in various strains (Supplementary Figure S7). ESBL-plasmid p2A_IncF carries a SPbeta-like prophage (68.4 kB), which encodes all 12 resistance genes of that plasmid. With the exception of p3A_crypt and pRE3B_crypt, all plasmids bigger than 35 kB carry plasmid addiction systems (22) (toxin-antitoxin (TA) systems, Supplementary Figure S8).

**Figure 1.**
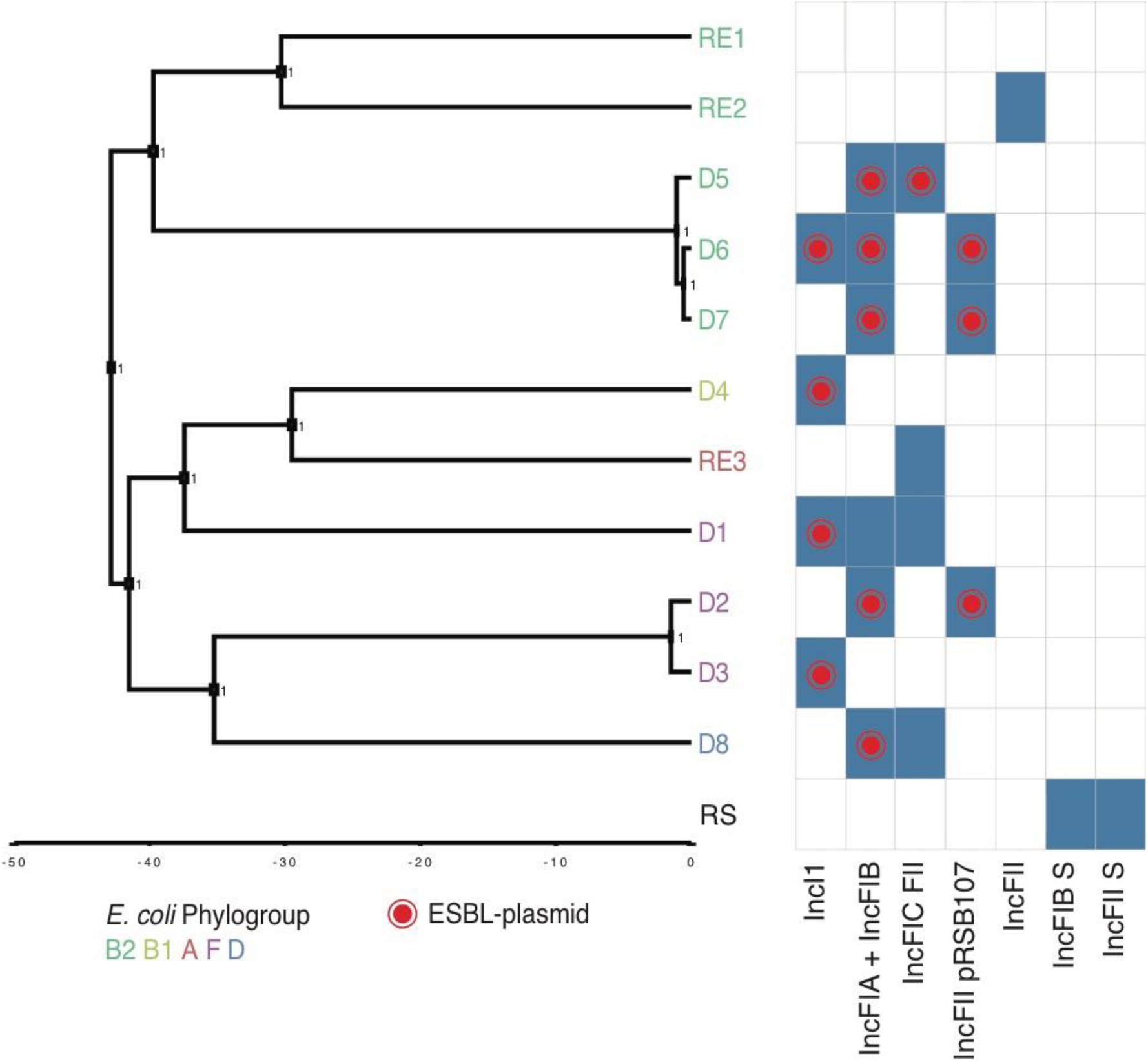
Phylogenetic tree of the *E. coli* donor and recipient strains, inferred using Bayesian inference on a core genome alignment. Strain names at the tips are coloured by *E. coli* phylogroup. The *S*. Typhimurium recipient RS was not included in the phylogeny, but listed here to allow comparison of the plasmid content. RE1-3 denotes the three *E. coli* recipients and D1-8 denotes the eight donors. Blue rectangles show that a plasmid with the indicated IncF or IncI replicon (incompatibility marker) is present in that strain. A red dot indicates the replicon(s) present on the ESBL-plasmid in each strain.

### Diverse clinical ESBL-plasmids spread extensively through recipient populations in the absence of antibiotics

To investigate the spread of ESBL-plasmids in the absence of antibiotics, we first performed conjugation experiments with all possible donor-recipient combinations (referred to as 1^st^ generation *in vitro* experiment). We use the final transconjugant frequency, i.e. the fraction of the recipient population that carried the ESBL-plasmid after 24 hours, to measure plasmid spread. The highest final transconjugant frequency (∼0.1%) was achieved when plasmid p4A_IncI spread in populations of recipient RE3. Five of the eight ESBL-plasmids spread in more than one of the *E. coli* recipients and their final transconjugant frequencies spanned 5 orders of magnitude (Figure 2A). The average final transconjugant frequency varied depending on the donor-plasmid pair and among recipient strains (two-way ANOVA excluding D2, D3, D7, effect of donor-plasmid pair: *F*_4,66_ *=* 87.665*, P* < 0.01, effect of recipient: *F*_2,66_ = 5.439, *P* < 0.01). The variation among donor-plasmid pairs depended also on the recipient (donor-plasmid pair×recipient interaction: *F*_8,66_ = 3.164, *P* < 0.01). Although these ESBL-plasmids are natively associated with *E. coli*, they reached comparable maximal transconjugant frequencies in the RS (i.e. *S*. Typhimurium) recipient populations (Figure 2B). The variation across donor-plasmid pairs was similar to that obtained with *E. coli* recipients, with the exception of p6A_IncI, which did not spread to recipient RS.

**Figure 2.**
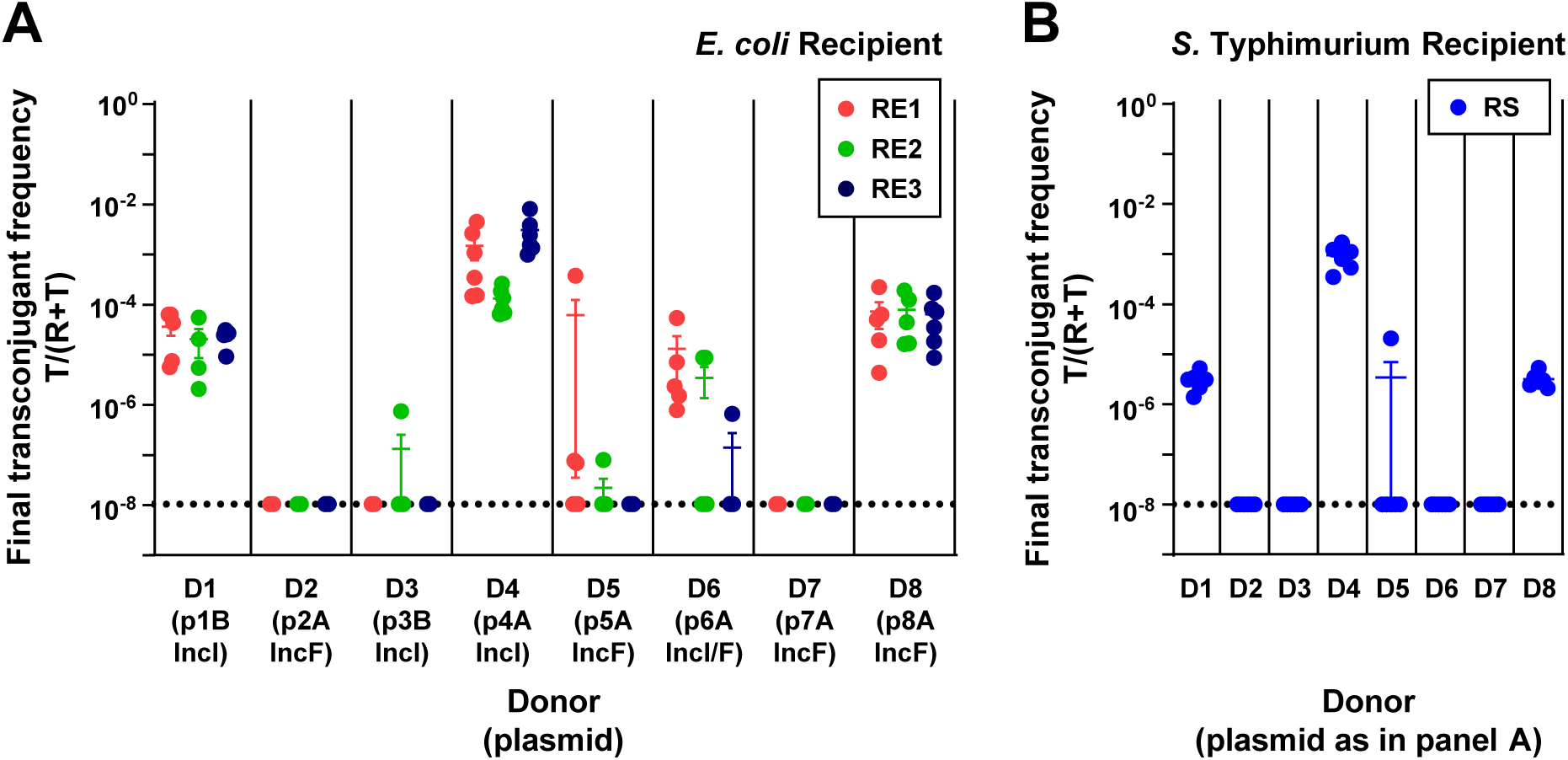
ESBL-plasmids spread at variable rates in the absence of antibiotics (1^st^ generation *in vitro* experiment). Plasmid spread was measured as the final transconjugant frequency, i.e. the ratio of the recipient population carrying the ESBL-plasmid (T), relative to the total of plasmid-free (R) and plasmid carrying (T) recipient populations. Final transconjugant frequency is shown for recipient populations of *E. coli* strains RE1-3 (A) and *S*. Typhimurium strain RS (B). Circles represent independent replicates (n=4-6) and the beams are mean values ± standard error of the mean (SEM). The detection limit was at ∼10^-8^. Total population densities can be found in Supplementary Figure S10.

The pili of type IncI and IncF plasmids support plasmid transfer on solid surface and in liquid growth environment (72). To investigate to which extent these ESBL-plasmids could transfer after plating on selective plates, we performed a surface-mating experiment for a subset of donor-recipient combinations (Supplementary Figure S9). On this solid surface, only p1B_IncI and p4A_IncI transferred from donor to recipient and resulted in transconjugant frequencies ranging from 10^-8^ to 10^-5^, in a recipient-dependent manner (two-way ANOVA excluding D6, D8 and RS: effect of recipient: *F*_2,18_ = 34.29, effect of donor-plasmid pair: *F*_1,18_ = 13.750, *P* < 0.01 in both cases). This suggests that most of the plasmid transfer in our conjugation experiments took place in the liquid growth environment.

### Plasmid, donor and recipient factors lead to variation in plasmid spread

In the 1^st^ generation *in vitro* experiment (Figure 2), each plasmid was present in a single donor and thus we could not separate the contributions of plasmid and donor strain to the observed plasmid spread. Therefore, we performed an second conjugation experiment (referred to as 2^nd^ generation *in vitro* experiment) under conditions identical to the 1^st^ generation *in vitro* experiment, but where each donor strain was represented with multiple different plasmids and each plasmid was represented in multiple donors. Specifically, for the 2^nd^ generation *in vitro* experiment, we used eight transconjugants isolated from the 1^st^ generation *in vitro* experiment as plasmid donors and three of the same recipient strains (Figure 3). The final transconjugant frequency varied among donor strains and among plasmids (three-way ANOVA with plasmid, excluding p1B_IncI, donor and recipient as factors, effect of donor strain: *F*_2,90_ = 150.133, *P* < 0.001, effect of plasmid: *F*_1,90_ = 49.717, *P* < 0.001). Variation among plasmids depended on both, the recipient and the donor strain (donor strain×plasmid interaction: *F*_2,90_ = 96.352, *P* < 0.001; recipient×plasmid interaction: *F*_2,90_ = 29.610, *P* < 0.001). For instance, when the donor and recipient strains were both RS, both IncI ESBL-plasmids yielded remarkably high final transconjugant frequencies of 40%. A second analysis supported variation among donor strains and plasmids and that variation among plasmids depended on recipient and donor strain (three-way ANOVA excluding RE2, effect of donor strain: *F*_1,93_ = 560.269, *P* < 0.001, effect of plasmid *F*_2,93_ = 156.075, *P* < 0.001, recipient×plasmid interaction: *F*_4,93_ = 26.104, *P* < 0.001, donor strain×plasmid interaction: *F*_2,93_ = 3.999, *P* = 0.022). As in our 1^st^ generation *in vitro* experiment, average final transconjugant frequencies also varied among recipients (*P* < 0.001 for effect of recipient in both three-way ANOVAs). Thus, the final frequency of transconjugants depended on donor strain, plasmid, and recipient. Also, we found plasmid spread to be especially efficient when donor and recipient are the same strain (Figure 3, self-self transfer).

**Figure 3.**
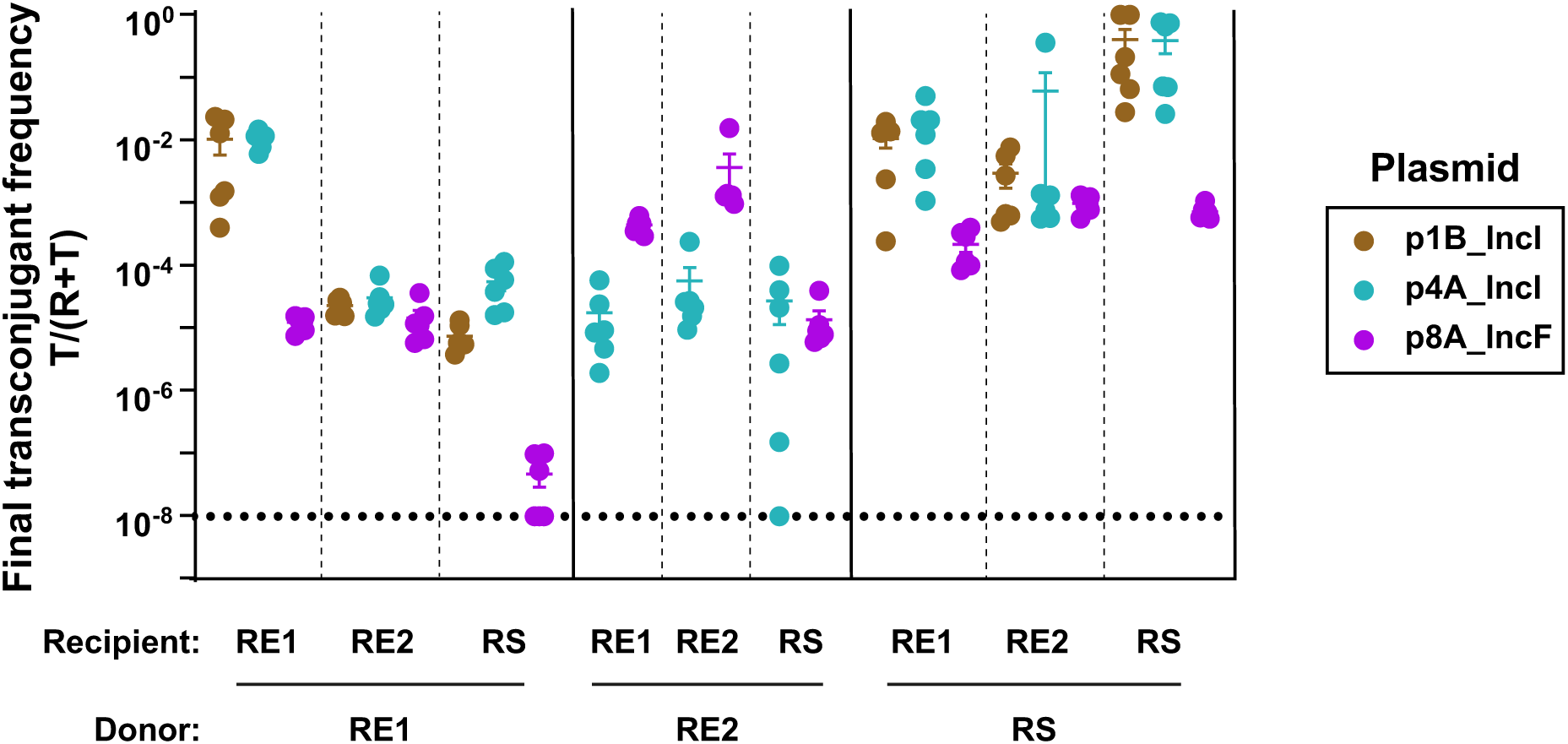
Final transconjugant frequency depends on donor, recipient, and plasmid (2^nd^ generation *in vitro* experiment). Eight transconjugants isolated from mating assays in the 1^st^ generation *in vitro* experiment (Figure 2), used here as plasmid donor strains, transferred their plasmid to three different recipients. Circles represent independent replicates (n = 6), the beams are mean values ± SEM and different plasmids are indicated in colour. The detection limit was at ∼10^-8^. Total population densities can be found in Supplementary Figure S11. Donor RE2 carrying plasmid p1B_IncI was excluded, see methods.

For some plasmid-recipient combinations we noticed that replacing the native donor strains with transconjugants from the 1^st^ generation *in vitro* assay (primary and secondary plasmid transfer, respectively) led to large differences in final transconjugant frequencies (Figures 2 and 3). For a subset of strains, we tested whether these resulted solely from the substitution of the native donor strain (D1) with a secondary plasmid host (RE1 and RS, Supplementary Figure S12). When RE1 and RS acted as a donor strain for plasmid p1B_IncI, transconjugant frequencies of RE1 carrying p1B_IncI increased 45-fold and 112-fold, respectively, compared to when p1B_IncI was transferred from its native donor strain D1. When both donor and recipient were RS, the final transconjugant frequency increased 2800-fold compared to transfer of plasmid p1B_IncI from its native host D1 to RS. Indeed, this showed that plasmid transfer from a secondary bacterial host can differ strongly from its transfer from the initial host.

### ESBL-plasmids can spread rapidly *in vivo*, with efficiencies correlating with the *in vitro* trends

Transfer of ESBL-plasmids in the human gut has only been confirmed by sequence-based studies (28, 73). However, to our knowledge the dynamics of their spread *in vitro* have not yet been thoroughly compared to any *in vivo* system. Therefore, we performed conjugation experiments over seven days in absence of antibiotics. These were done in gnotobiotic mice with a defined microbiota (46), with three clinical donors (D4, D8, and D7), and two recipients (RE2 and RE3, Figure 4). The residential microbiota allows colonization of approximately 10^8^ *E. coli* per gram faeces, *E. coli* densities representative of the guts of some humans and animals (42, 74). The variation of ESBL-plasmid spread *in vivo* was in qualitative agreement with the 1^st^ generation *in vitro* experiment. As in the 1^st^ generation *in vitro* experiment (Figure 2) for recipient RE3 we observed highest transconjugant frequencies with plasmid p4A_IncI followed by p8A_IncF and no transconjugants with p7A_IncF (Figure 4). Furthermore, the final transconjugant frequency with plasmid p4A_IncI was higher with RE3 than RE2 both *in vivo* and *in vitro* (see Figure 4A-B, blue dots, and Figure 2). Lastly, the final transconjugant frequencies with p8A_IncF were similar for both recipient populations (Figure 4A-B, purple dots, and Figure 2). Because the final frequency of RE3 carrying p4A_IncI (1%) was already reached at day 1, we re-performed this conjugation experiment, sampling more densely in time and found the final transconjugant frequency to be established already eight hours after the orogastric introduction of donor D4 (Supplementary Figure S13). This rapid increase in transconjugant frequency was followed by a 6-day plateau, which may result from the simultaneous decrease of recipient and transconjugant populations over time (Supplementary Figure S14). Indeed, direct competition experiments *in vivo* (Supplementary Figure S15) confirmed the competitive advantage of donor D4 over RE3. This fitness benefit can perhaps be explained by the difference in *in vitro* growth rate estimates (Supplementary Figure S16).

**Figure 4.**
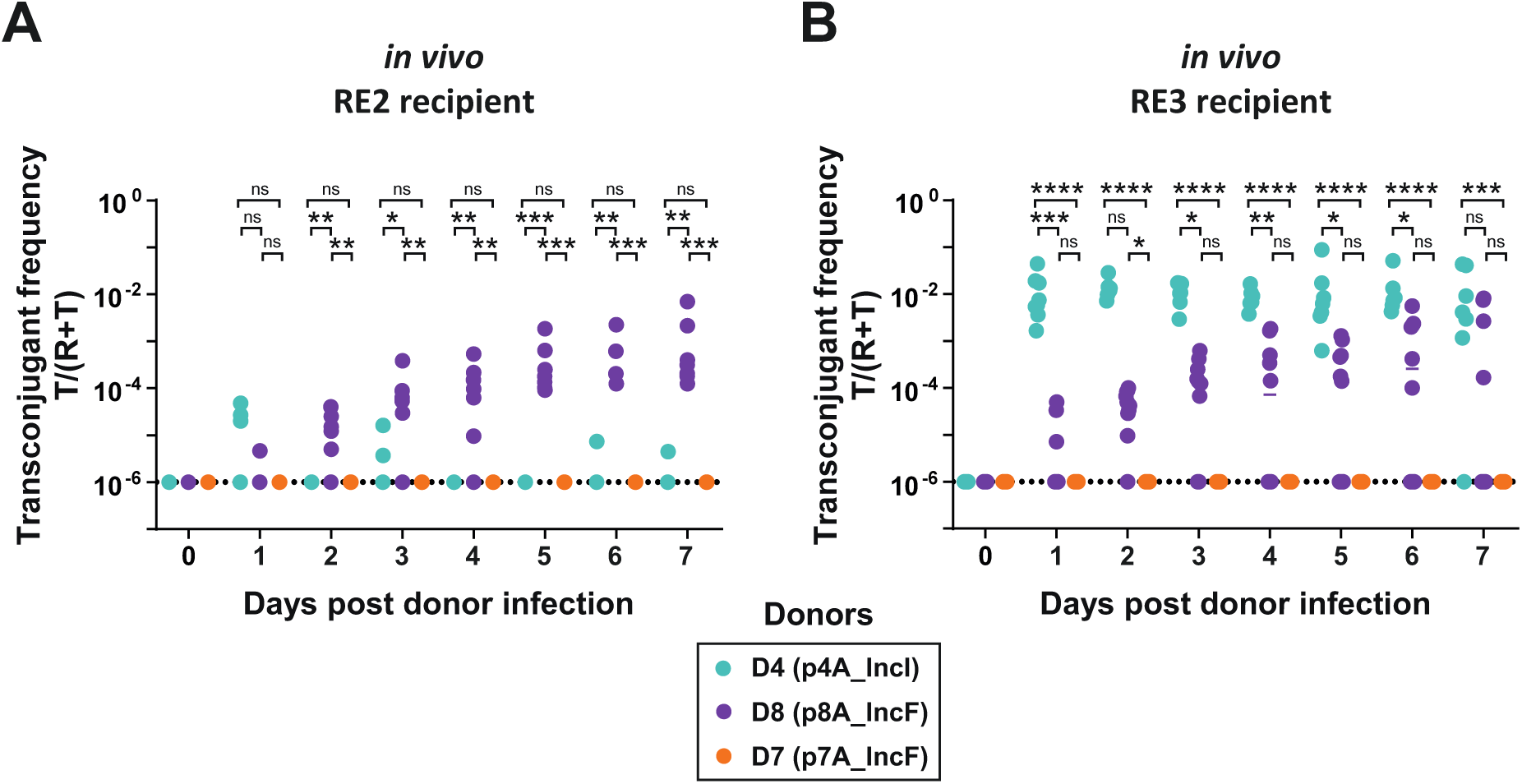
ESBL-plasmids can spread in the gut in the absence of antibiotic selection. We measured the spread of three plasmids as final transconjugant frequency in two distinct recipient populations, A) RE2 and B) RE3, and enumerated transconjugants in faeces by selective plating. Dotted lines indicate the detection limit for selective plating. Circles represent independent replicates (n = 7 for RE2 conjugations; n =7 for D4-RE3; n = 10 for D8-RE3 and D7-RE3), lines show the median and different donor-plasmid pairs are indicated in colour. Kruskal-Wallis test p>0.05 (ns), p<0.05 (*), p<0.01 (**), p<0.001 (***), p<0.0001 (****). Total population densities can be found in Supplementary Figure S14.

### ESBL-plasmids generate no significant cost after transfer to new bacterial hosts *in vitro* and *in vivo*

We investigated the effect of plasmids on bacterial growth in the absence of antibiotics for ten strain-plasmid combinations *in vitro*. We estimated plasmid cost as the growth rate of transconjugants relative to their respective plasmid-free recipient strain and found no significant effect of plasmid carriage (Figure 5A-B, Student’s t-Test for *E. coli* hosts and Wilcoxon Rank Sum Test for *S.* Typhimurium, *P* > 0.05 in all cases, before and after Holm’s correction for multiple testing). *In vivo*, we investigated the effect of ESBL-plasmids p4A_IncI and p8A_IncF on bacterial fitness with direct 1:1 competition between recipient RE3 and its transconjugants (Figure 5C). After seven days of competition, for transconjugants with p4A_IncI, there was no significant change in the relative frequency of recipients to transconjugants. Transconjugants carrying p8A_IncF outcompeted RE3 leading to a relative frequency of ∼0.1 after seven days, indicating a minor fitness advantage by carriage of p8A_IncF.

**Figure 5.**
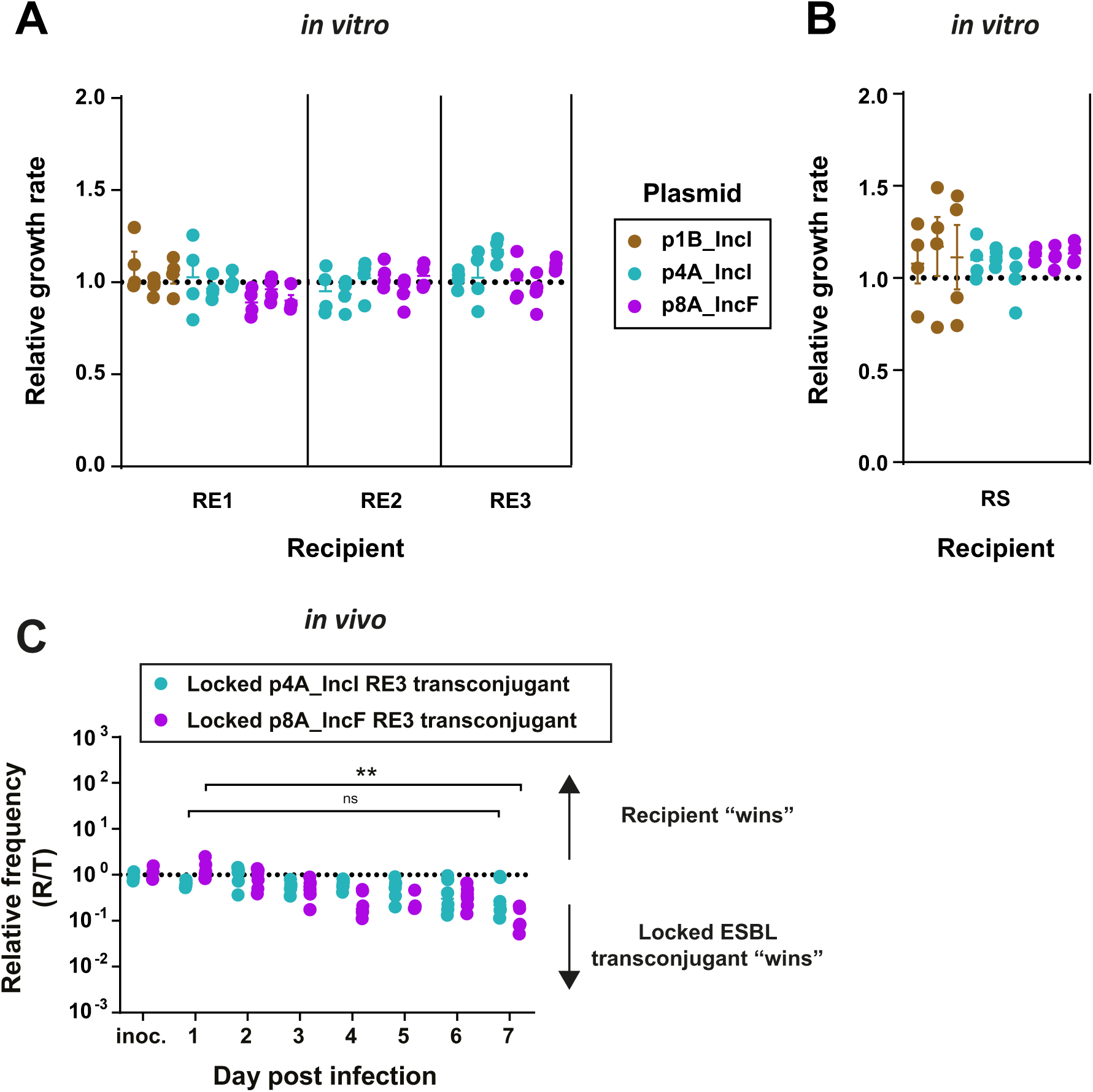
No evidence for cost of plasmid carriage for transconjugants. A-B) We measured plasmid cost for ten strain-plasmid combinations, with three independently isolated transconjugants each (n = 4; beams are mean values ± SEM). Transconjugants and their plasmid free complements grew in independent cultures and we calculated the relative growth by dividing the transconjugant growth rates (h^-1^) by the mean growth-rate of plasmid-free strains. C) We performed the competition experiment by colonizing the mice with a 1:1 mix of a non-conjugative transconjugant (*oriT*-knockout) and recipient RE3 (n = 6; 3 independent transconjugants, n = 2 for each). (C) Kruskal-Wallis test p>0.05 (ns), p<0.01 (**).

### Plasmid transfer rates drive plasmid spread and their *in vitro* estimates are a good proxy for *in vivo* transfer

The observed variation in final transconjugant frequencies *in vitro* and the dynamics in the mouse gut could be driven by horizontal plasmid transfer or by clonal expansion of transconjugants. Although non-significant, *in vitro,* the *S*. Typhimurium transconjugants showed a consistent trend towards higher relative growth rates (Figure 5B). Our calculations demonstrated that differences in growth rates resulting from ESBL-plasmid carriage could indeed explain final transconjugant frequencies of *S.* Typhimurium recipients carrying p1B_IncI and p8A_IncF, but not plasmid spread in any *E. coli* recipient population (Supplementary Results). To estimate plasmid transfer rates, we applied the Simonsen method to population densities obtained in the 1^st^ generation *in vitro* experiment (S3 Table) (69). We found that conjugative transfer rates of primary transfer, from donor to recipient, correlate strongly with observed final transconjugant frequencies, across all donor-recipient combinations (Supplementary Figure S17, Pearson’s test, *r =* 0.99, *P* < 0.001). Together, this suggests that horizontal plasmid transfer from donor to recipient dictated observed spread of ESBL-plasmids *in vitro*.

The *in vivo* competition experiment revealed a growth advantage of RE3 when carrying p8A_IncF, allowing a two-fold increase of the initial transconjugant frequency when growing for seven days in the gut (from 0.5 to 0.9, Figure 5C). Because we used *oriT*-mutants, no plasmid could be horizontally transferred during this competition experiment and therefore increasing transconjugant populations must have resulted from clonal growth. Allowing for horizontal plasmid transfer (seven-day conjugation experiment), however, the transconjugant frequency of RE2 and RE3 carrying p8A_IncF increased from the detection limit of 10^-6^ up to final frequencies of 1% (Figure 4A-B). This large difference in transconjugant population increase with and without conjugation allows us to conclude that in our gut colonization model without antibiotic selection, the spread of ESBL-plasmids was driven by conjugative transfer, rather than by clonal expansion of transconjugants. The fast increase of the transconjugant population RE3 carrying p4A_IncI within only eight hours (Supplementary Figure S13), despite a lack of growth advantage over recipient RE3 (Figure 5C), further supports this result. *In vitro*, we found that transfer from donor strains and from transconjugants to recipient strains could vary several orders of magnitude (Supplementary Figure S12). Such a difference in primary and secondary plasmid transfer could also be present *in vivo*. However, the transconjugant populations were minor compared to the size of the donor populations throughout the *in vivo* experiment (Supplementary Figure S14). Thus, we can conclude that most transfer events were between donors and recipients, rather than between transconjugants and recipients. Altogether, we showed horizontal transfer to allow for rapid ESBL-plasmid spread in the murine gut, in the absence of antibiotic selection, and demonstrated that our *in vitro* transfer rates reflect *in vivo* transfer dynamics.

### Plasmid factors are the main determinant of observed plasmid transfer rates

We demonstrated that the extent of the spread of ESBL-plasmids in both *in vitro* and *in vivo* systems was a result of their transfer rates. To investigate genetic factors that might influence plasmid transfer rates, we performed detailed bioinformatic analyses on plasmids, donor and recipient genomes. We found that the presence of essential *tra* genes on ESBL-plasmids is the main genomic factor predicting their spread (Figure 6): the 5/8 ESBL-plasmids encoding all necessary *tra* genes spread to at least three of the five recipient strains.

**Figure 6.**
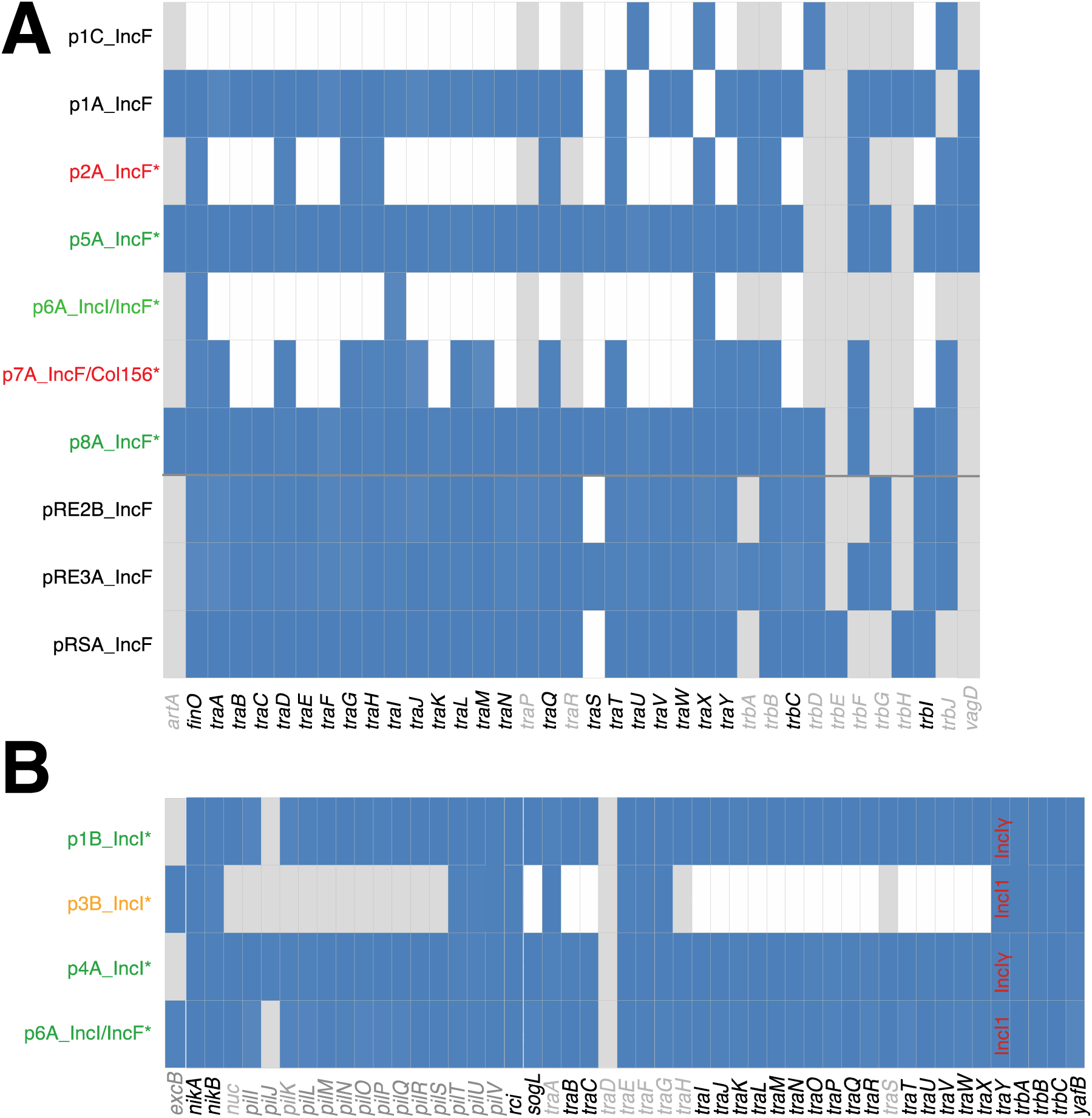
Transfer genes. Presence (blue) or absence (white) of essential *tra* genes for IncF plasmids (A), and IncI plasmids (B). Genes and panels in grey are non-essential for pilus biogenesis, DNA transfer or conjugation, according to Koraiman (39) (A) and Komano (37, 38) (B). ESBL-plasmids are indicated by an asterisk: green indicates spread to multiple recipient populations, orange indicates conjugation with only one replicate population, and red indicates no transfer. Non-ESBL plasmids are labelled in black. Plasmid p6A_IncI/IncF is shown on both panels A and B, but only the IncI1 transfer system is complete.

Our experiments showed that the more fine-scale variation in plasmid spread depends on donor and recipient factors (Figures 2-3). We investigated whether the phylogenetic relatedness of strains (Figure 1) could explain observed plasmid transfer dynamics. Qualitatively, the relatedness of mating donor and recipient strains was not correlated with transfer rates: D5 and D6 are equally closely related to recipients RE1 and RE2, yet we observed very different final transconjugants frequencies (Figure 2). Additionally, recipient RS is phylogenetically distant to all the donors, yet we observed variation in final transconjugants frequency similar to those observed across *E. coli* recipients. In contrast, the relatedness of plasmids with regard to their replicons, because plasmids carrying the same replicon can pose a barrier to each other, seemed to be a good predictor of plasmid spread. The final transconjugant frequencies of the IncFII ESBL-plasmids p5A_IncF and p6A_IncI/F varied largely depending on recipients (Figure 2). They were highest with recipient RE1, likely because it is the only recipient without a plasmid encoding an IncFII-replicon. The lack of transconjugants resulting from conjugation of donor D6 carrying plasmid p6A_IncI/F with recipient RS, could be due to the incompatibility with the resident plasmid pRS_IncF. This is intriguing because the presence of *tra* genes (Figure 6) indicates that plasmid p6A_IncI/F transfers with the IncI1 transfer machinery, whereas the incompatibility can only be IncFII based.

We sequenced transconjugants resulting from both *in vitro* and the *in vivo* conjugation experiments (Supplementary Table S2). This provided further evidence for the role of incompatibility. Plasmid p8A_IncF carries half of the IncFIC-FII replicon (Supplementary Figure S5) and resulted in the loss of the resident F-plasmid pRE3A_IncF in recipient RE3, both *in vitro* and *in vivo*. This plasmid interference, however, seemed not to limit the transfer of plasmid p8A_IncF into RE3, because it spread to all recipients at the same rate (Figure 2).

Donor and recipient strains carry multiple non-ESBL plasmids that could have been transferred during conjugation experiments. Sequencing revealed only a few isolated cases of co-transferring plasmids *in vitro* and none *in vivo*. For plasmid p8C_IncBOKZ we found some transfer to all four recipients of the 1^st^ generation experiment and from there even to further recipients in the 2^nd^ generation experiment. This process seemed independent of donor and recipient strains (Supplementary Table S2). The only small Col-plasmids that were transferred were p1D_ColRNAI and p8G_Col8282. The P1 phage-like plasmid p8B_p0111 transferred *in vitro* to RE3 but not *in vivo* (Supplementary Table S2). Although they encode the *tra* genes (S2 File), the resident plasmids in the recipients (RE2, RE3 and RS) showed no horizontal transfer. Sequencing transconjugants also allowed us to screen for mutations that could explain some of the variation in final transconjugant frequencies across donor-recipient pairs or between primary and secondary plasmid transfer (Figures 2-3). We found no mutations on ESBL-plasmids, neither after the 24-hours *in vitro* nor the 7-days *in vivo* conjugation experiments (Supplementary Table S2). The only consistent changes on plasmids were the mutations of P1 phage-like plasmid p8B_p0111, which was passed to recipient RE3 *in vitro,* but not *in vivo*: here we found mutations in the genomic region encoding side tail fibre proteins. Similar mutations were also frequently present within chromosomal prophages. We found several other chromosomal mutations in transconjugants (Supplementary Table S2). For instance, all transconjugants resulting from conjugation into RE1 recipients (*n = 11*) and some (*n = 9* out of 22) RE3 recipients showed intergenic mutations in the promoter sequence or the phase ON/OFF region of the *fim* operon. The *fim* genes encode the type 1 pilus (type 1 fimbria), a virulence factor responsible for cell adherence (75). RE1 chromosomally encodes for two mobilization proteins mobA, required for the mobilization of plasmid and conjugative transposons. In all 11 sequenced RE1-transconjugants, we found various intergenic mutations upstream of at least one *mobA*.

We found various RM systems (Supplementary Figure S3) and investigated whether ESBL-plasmid transfer from an RM deficient donor into a recipient with this RM system could explain reduced plasmid transfer rates. We found high rates of self-self transfer (Figure 3), which could result from identical RM systems in donor and recipient strains but for other mating pairs, we found no relation between RM systems and transfer rates. The anti-restriction protein YfjX (ardB) is present in nearly all strains, including on all ESBL-plasmids, and could reduce the detrimental effect of RM systems in recipients. We found the adaptive immunity systems CRISPR-Cas Type 1F (recipients RE1 and RE2) and CRISPR-Cas Type 1E (recipients RE3, RS and donors D1-4, D8). We investigated whether they could be a barrier to conjugation, by screening whether the spacer sequences in recipient strains matched any of our plasmids or phages. This was not the case. We further investigated the activity of the CRISPR system *in vitro* and *in vivo* by screening all CRISPR arrays in sequenced transconjugants for spacer sequences that were not present in the recipients prior to conjugation. We did not observe spacer acquisition, neither *in vitro* nor *in vivo*.

## Discussion

Conjugative plasmids are a key driver of the antibiotic resistance crisis through their ability to spread within and between bacterial species. Many possible factors affecting their spread have been identified, but most studies investigate these factors *in vitro*, in laboratory strains or in isolation (13–15, 27, 76). Understanding the relevance of these factors in ecologically complex environments, where plasmid transfer typically occurs, necessitates studies that aim to untangle their relative contributions. Here we combined bioinformatic analyses with *in vitro* and *in vivo* experiments and provide a comprehensive and quantitative investigation of plasmid spread in the absence of antibiotic selection.

We found that differences in horizontal plasmid transfer and not cost of plasmid carriage are the main source of variation in plasmid spread. In this study, ESBL-plasmids encoding the essential genes for conjugation spread efficiently and the variation of their spread depended also on recipient and donor strain. This is of particular importance because approximately half of all known plasmids can transfer via conjugation (16) and thus, a substantial amount of antibiotic resistance plasmids have the potential to spread through diverse bacterial populations without the need for strong selective forces, such as antibiotics.

Our measurements of plasmid spread *in vitro* showed good qualitative agreement with plasmid spread in a mouse model, which was reflected in the ranking of final transconjugant frequencies and its dependence on the recipient strain. This suggests that key aspects of plasmid spread in complex environments can be quantified with higher throughput *in vitro* testing. *In vivo*, the rise of the transconjugant frequency was remarkably fast. RE3 carrying plasmid p4A_IncI reached a frequency of 1% after 8 hours, but surprisingly plateaued thereafter. Explanations thereof could be the outcompetition of recipient and transconjugant populations by the plasmid donor strain or a reduced growth rate after initial colonization of the plasmid donor strain. The latter one, however, seems unlikely as we would expect this to lead to a similar pattern for D8. Alternatively, this could result from spatial heterogeneities of bacterial populations. A high initial plasmid transfer rate with a plateau after one day has previously been observed for biofilm assays and in a streptomycin-treated mouse model (77) and was attributed to the fixed spatial position of donor and recipient cells in the biofilm and the mucus layer of the gut, respectively. We found that on a solid surface, p4A_IncI spread at 100-fold higher rates in RE3 than RE2 recipient populations, suggesting that plasmid transfer with recipient RE3 could benefit from a structured environment. In the colonization model used for our conjugation experiments (45), such interactions of bacteria with the host intestinal lining could also allow for more structured populations (78), and may indeed result in the observed transfer dynamics of plasmid p4A_IncI.

In natural systems bacteria often harbour multiple plasmids, which affect their ability to acquire further plasmids. Using bioinformatic analyses, we studied various plasmid features and related those to observed plasmid spread. We found plasmid incompatibility to play a role in conjugation with ST131 strains D5 and D6, but also to be a permeable barrier. This might be explained by the multiple replicons encoded on ESBL-plasmids, a strategy to circumvent limitations in spread due to plasmid incompatibility (79). Co-transfer of other plasmids has also been proposed to affect plasmid transfer rates (80), but we found little plasmid co-transfer and could not relate its occurrence to observed plasmid spread. Sequencing of the ESBL-plasmids in transconjugants revealed no mutations after transfer. This is in agreement with earlier findings reporting the absence of mutations on ESBL-plasmids even after 112 days of evolution of transconjugants (81). This suggests that clinical ESBL-plasmids are well adapted to Enterobacteriaceae and do not require clone-specific adaptations for successful spread.

Using bioinformatic analysis we also studied how strain-specific features influence plasmid spread. We found that all four recipient strains encode the adaptive immunity system CRISPR-Cas Type I, of which Type IF (recipients RE1 and RE2) is commonly associated with antimicrobial susceptibility in *E. coli* (82). It is generally believed that in laboratory *E. coli* strains such as K12, Type I CRISPR-Cas loci are inactive under laboratory growth conditions (83). This was also suggested by our investigation of spacer acquisition. Other bacterial defence systems are also known to affect the efficiency of plasmid transfer (84). For instance, it has been proposed that plasmid transfer to close kin is more efficient due to the similarity in RM systems of donor and recipient strains (27). Here, however, we did not find this relation. The ESBL-plasmids used in this study employ anti-restriction strategies like many other conjugative plasmids (85), and thus, we and others suggest that RM systems may only marginally shape horizontal plasmid transfer in natural systems (86, 87).

Given the central role of plasmids in the spread of antibiotic resistance, it is of great importance to understand the factors contributing to plasmid spread under natural conditions. Studies focusing on well-defined plasmid and laboratory strains have identified multiple factors relevant for plasmid spread under laboratory conditions. How these factors play together to affect plasmid spread under natural conditions, is key to understand why clinically relevant combinations of bacterial strains and plasmids, such as *E. coli* ST131 with IncFII plasmids encoding *bla*_CTX-M_, have become global threats (4–6). Encouragingly, the qualitative agreement between the *in vitro* measurements and the plasmid spread in the mouse model, suggests that *in vitro* testing with relevant clinical strains and plasmids may potentially be a good proxy for plasmid spread also in humans. Linking large-scale *in vitro* conjugation experiments using clinically relevant plasmid-strain combinations with epidemiological information on their spread thus seems an important avenue of future research to delineate the factors contributing to the global spread of antibiotic resistance by plasmids.

## Acknowledgements

We would like to acknowledge the members of the Bonhoeffer, Hall, Hardt, Diard, Egli, Stadler, and Ackermann labs, as well as Sebastien Wielgoss, for helpful discussions. We would also like to thank the staff at the RCHCI and EPIC animal facilities at ETH Zurich, particularly Sven Nowok; as well as the NGS team and Dr. Daniel Wuethrich of the Clinical Bacteriology and Mycology department at the University Hospital Basel for their excellent technical support. This work was funded by the NRP72 SNF grant (407240–167121) to SB, AH, AE, MA, TS and WDH. Additionally, AE, WDH and SB were supported by the Gebert Rüf Foundation “Microbials” programme grant “displacing ESBL”, EB by the Boehringer Ingelheim Fonds PhD Fellowship and MD by the SNSF professorship grant PP00PP_176954.

## Supplementary Figures, Tables and Results

**Figure S1.**
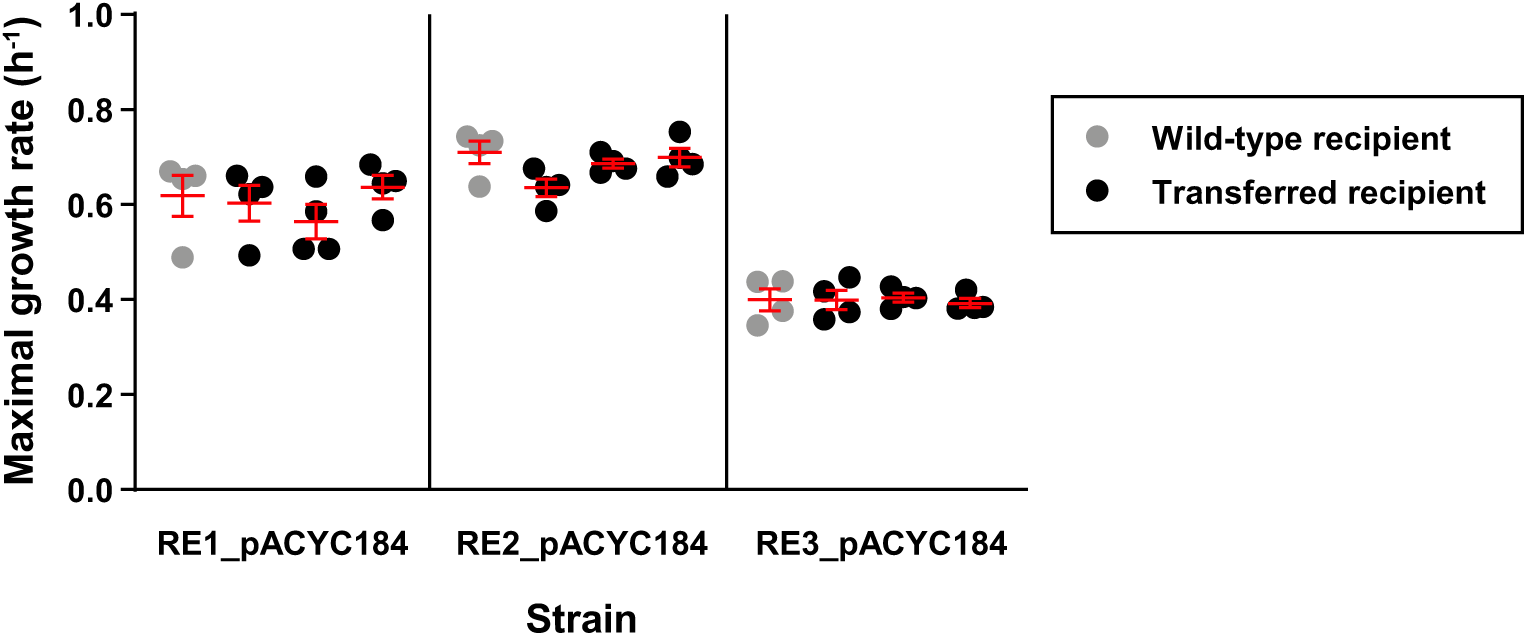
Control for transconjugant growth rates. We exposed recipients (RE1-RE3, *n = 4*) to the same culture handling as a conjugation assay would (black circles) to test for possible growth rate differences of recipients and transconjugants (Figure 5A-B) due to growth during the 1^st^ generation conjugation experiment (see Materials and methods). Beams are mean values ± SEM.

**Figure S2.**
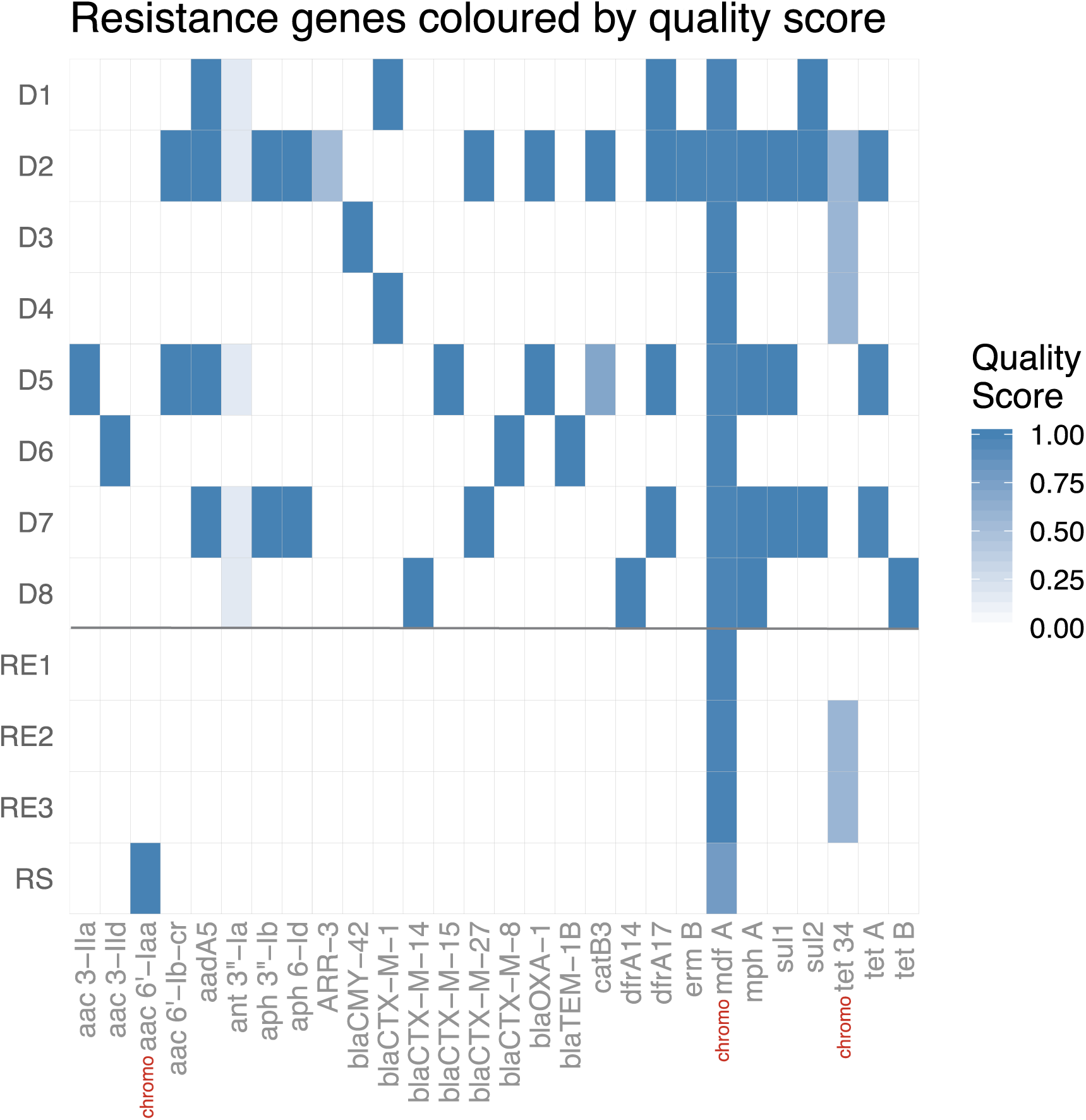
Resistance genes. Presence (blue) or absence (white) of resistance genes in donors (D1-D8) and recipients (RE1-RE3, RS). The quality score *q* reflects the percent identity *p* and coverage *c* of the BLAST match to the listed genes, where 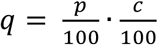. The genes ant 3”-Ia and ARR-3 were excluded from the S1 table. Chromosomal resistance genes are indicated with the red abbreviation chromo.

**Figure S3.**
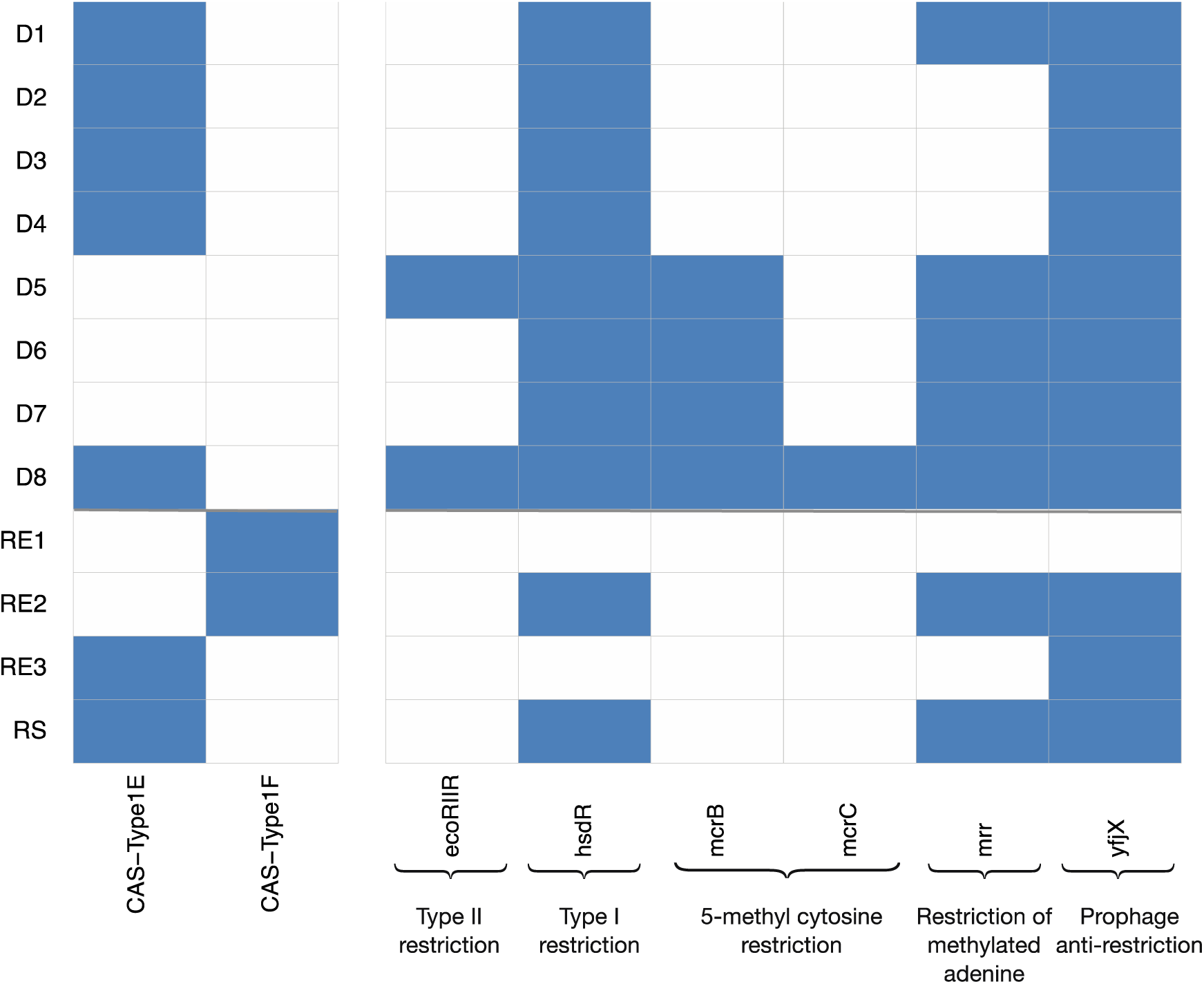
Bacterial immunity systems. Presence (blue) or absence (white) of CRISPR-Cas and restriction-modification systems (RM) in donors (D1-D8) and recipients (RE1-RE3, RS). CRISPR-Cas and RM systems were determined using prokka and confirmed using Rebase.

**Figure S4.**
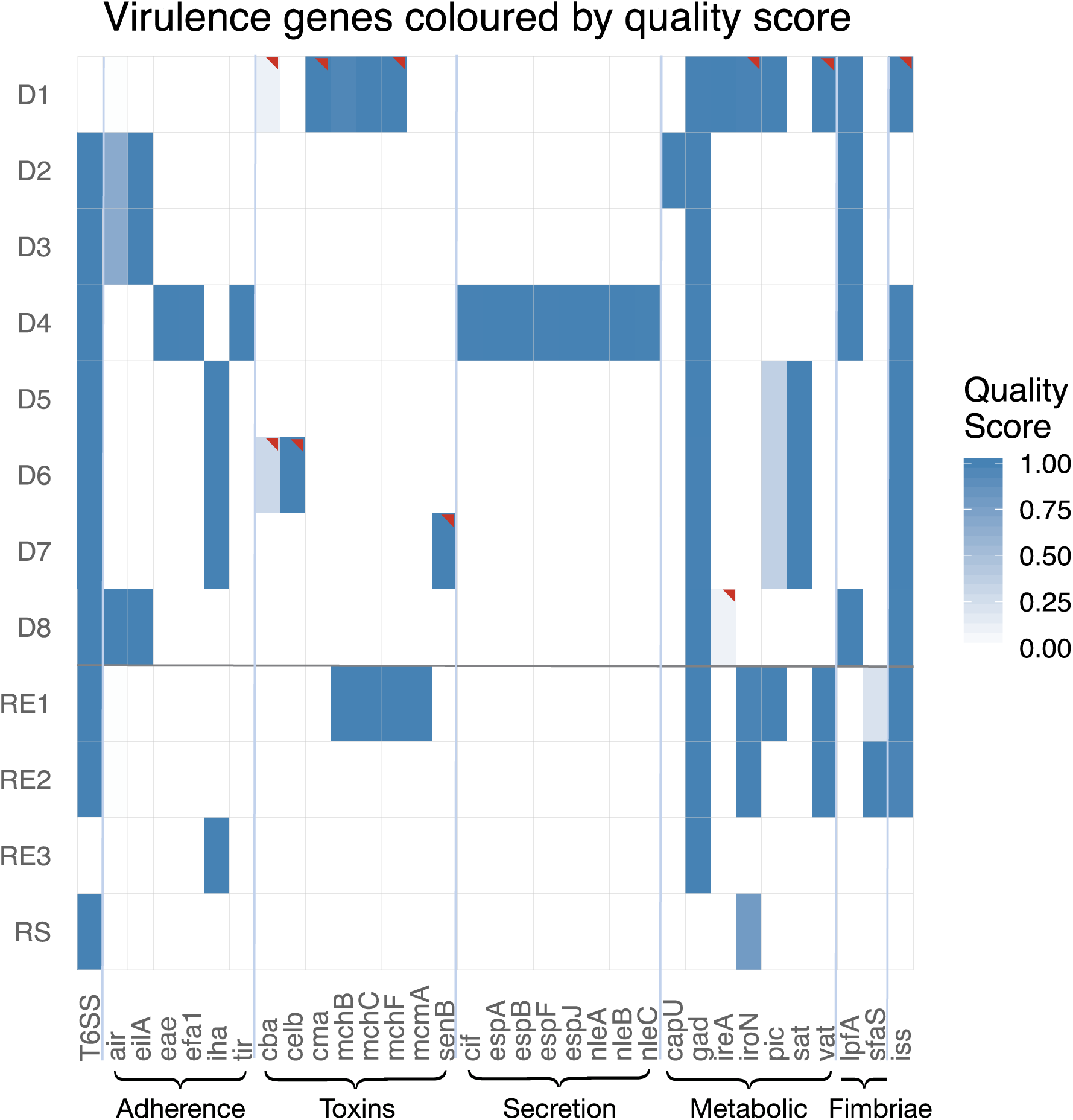
Type 6 secretion systems and virulence genes. Presence (blue) or absence (white) of Type 6 secretion systems (T6SS, defined by the presence of more than one prodigal T6SS-encoding gene in an operon) and virulence genes (found by BLAST) in donors (D1-D8) and recipients (RE1-RE3, RS). The quality score *q* reflects the percent identity *p* and coverage *c* of the BLAST match to the listed genes, where 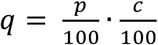. Plasmid-based sequences are flagged with a red triangle. The only strain with a pathogenic virulence profile is D4, which carries chromosomally encoded intimin genes (*eae*, *tir*) commonly associated with Enteropathogenic *E. coli*. All strains except RE3 and D1, encode for a T6SS.

**Figure S5.**
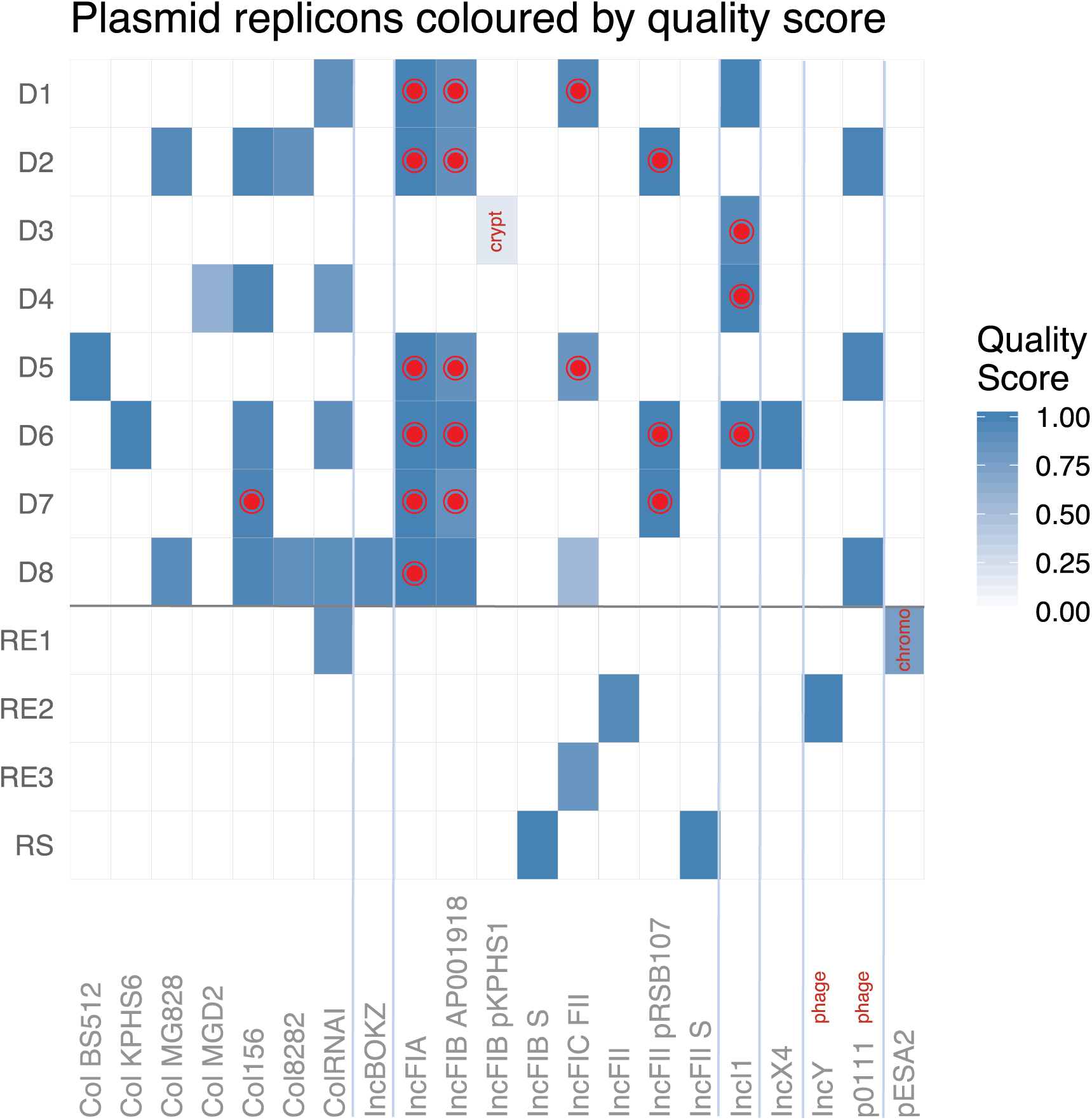
Plasmid replicons. Presence (blue) or absence (white) of plasmid replicons in donors (D1-D8) and recipients (RE1-RE3, RS). A red dot indicates the ESBL-plasmid, which can carry multiple replicons. The quality score *q* reflects the percent identity *p* and coverage *c* of the BLAST match to the listed genes, where 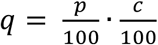. The matching regions of p3A_crypt and p8A_IncF were too short (< 75% coverage) to be included into S1 table. IncY and p0111 are P1-like phages.

**Figure S6.**
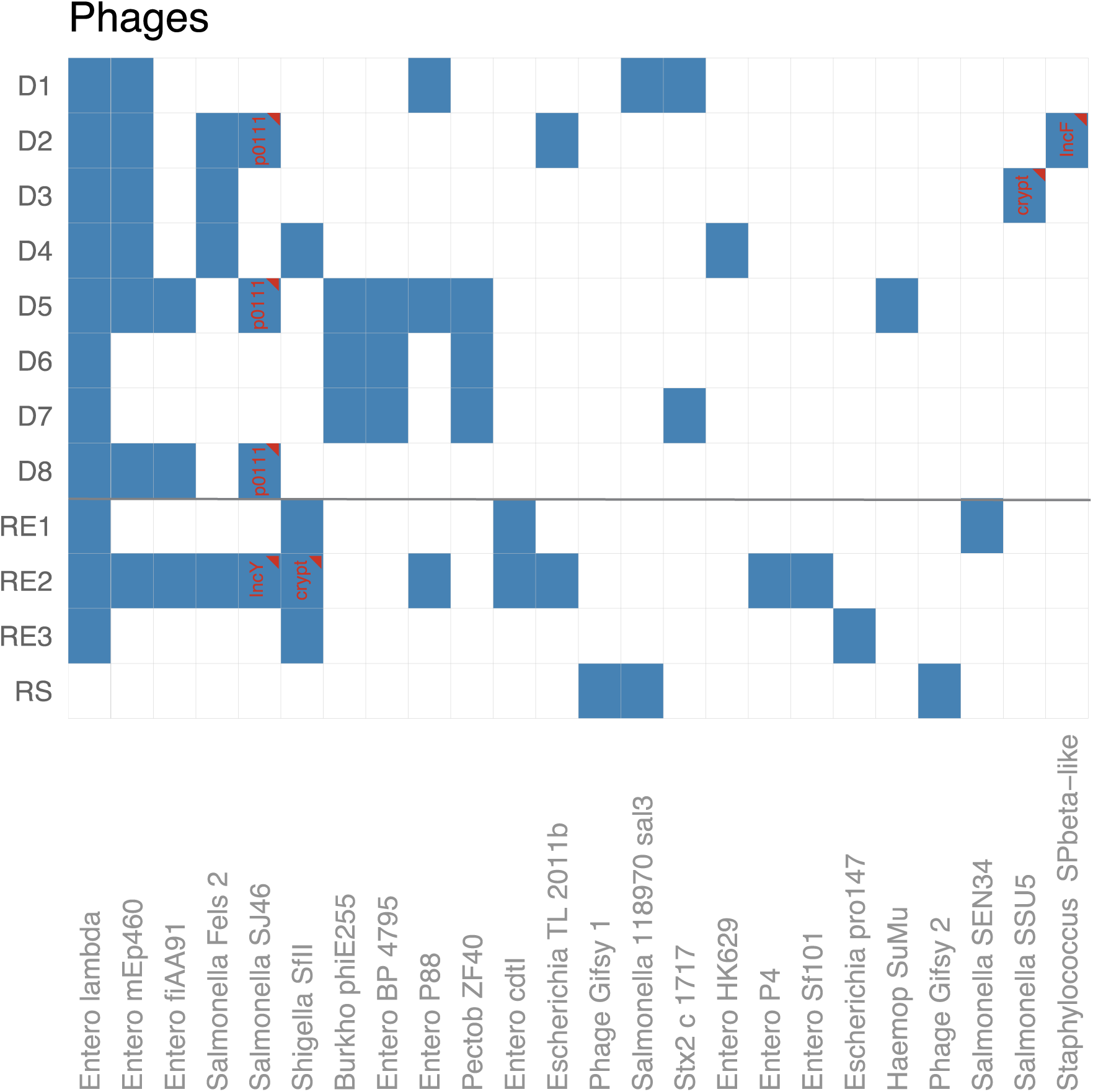
Phages. Presence (blue) and absence (white) of (pro)phages in donors (D1-D8) and recipients (RE1-RE3, RS). Plasmid-like phage sequences are flagged with a triangle and the replicon of the contig they were found on, both in red. All strains contain extensive (pro)phage related sequences.

**Figure S7.**
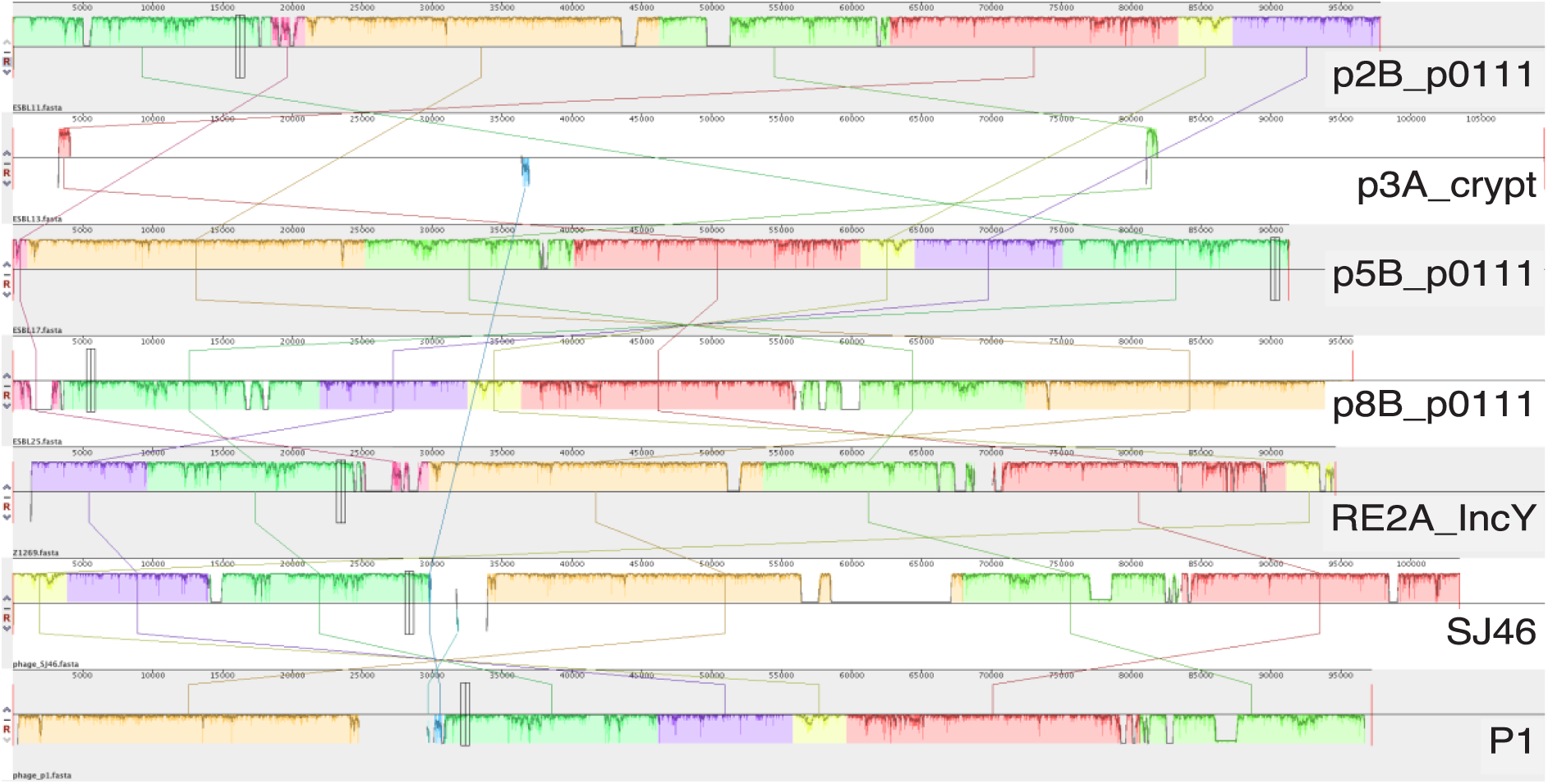
Mauve alignment. This alignment shows that the p0111 “plasmids” of D2, D5, D8, and the IncY “plasmid” of RE2 are highly related to each other and to the phages SJ46 and P1. Although highly similar to the phage SSU5, the p3A_crypt “plasmid” of D3 was largely unrelated to the P1-like phages.

**Figure S8.**
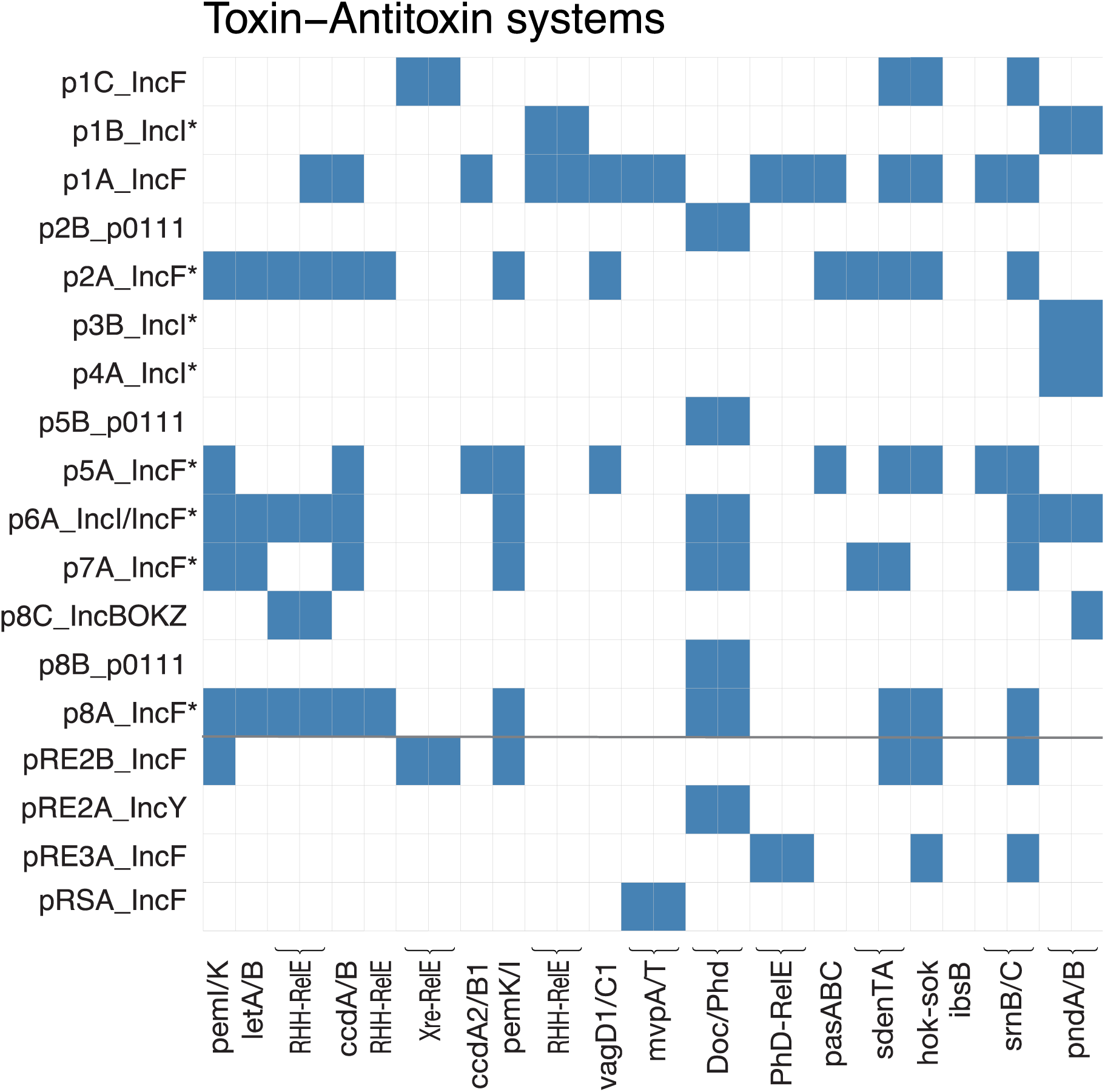
Toxin/Antitoxin-systems. Presence (blue) or absence (white) of Toxin/Antitoxin-systems (TA) in donors (D1-D8) and recipients (RE1-RE3, RS). Brackets indicate T/AT-pairs. The ESBL-plasmids are indicated by an asterisk. IncI1-plasmids and IncF-plasmids are generally associated with distinct TA-systems.

**Figure S9.**
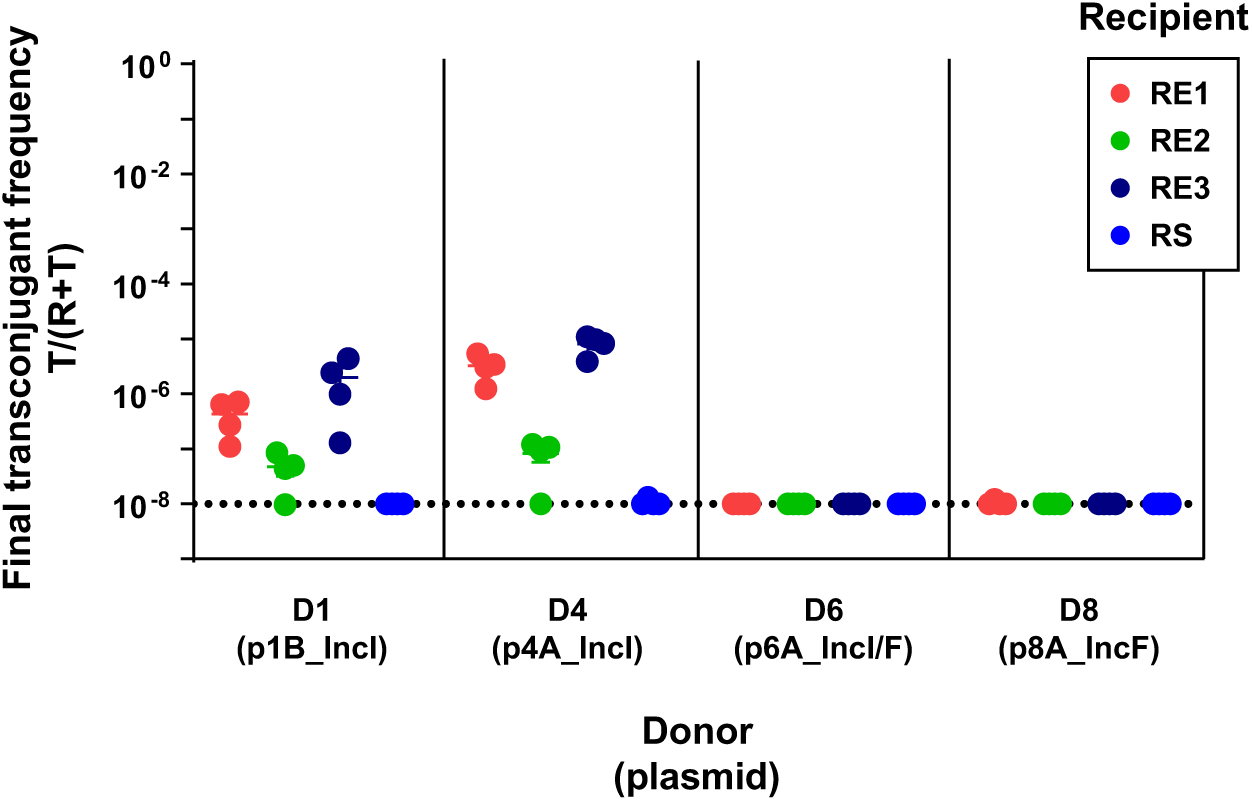
Surface-mating experiment. We allowed D1, D4, D6 and D8 to conjugate with all four recipients on LB-agar plates (*n = 4*). Resulting transconjugant frequencies depended on donor-plasmid pair and the recipient strain. Beams are mean values ± SEM, the dotted line indicates the detection limit by selective plating.

**Figure S10.**
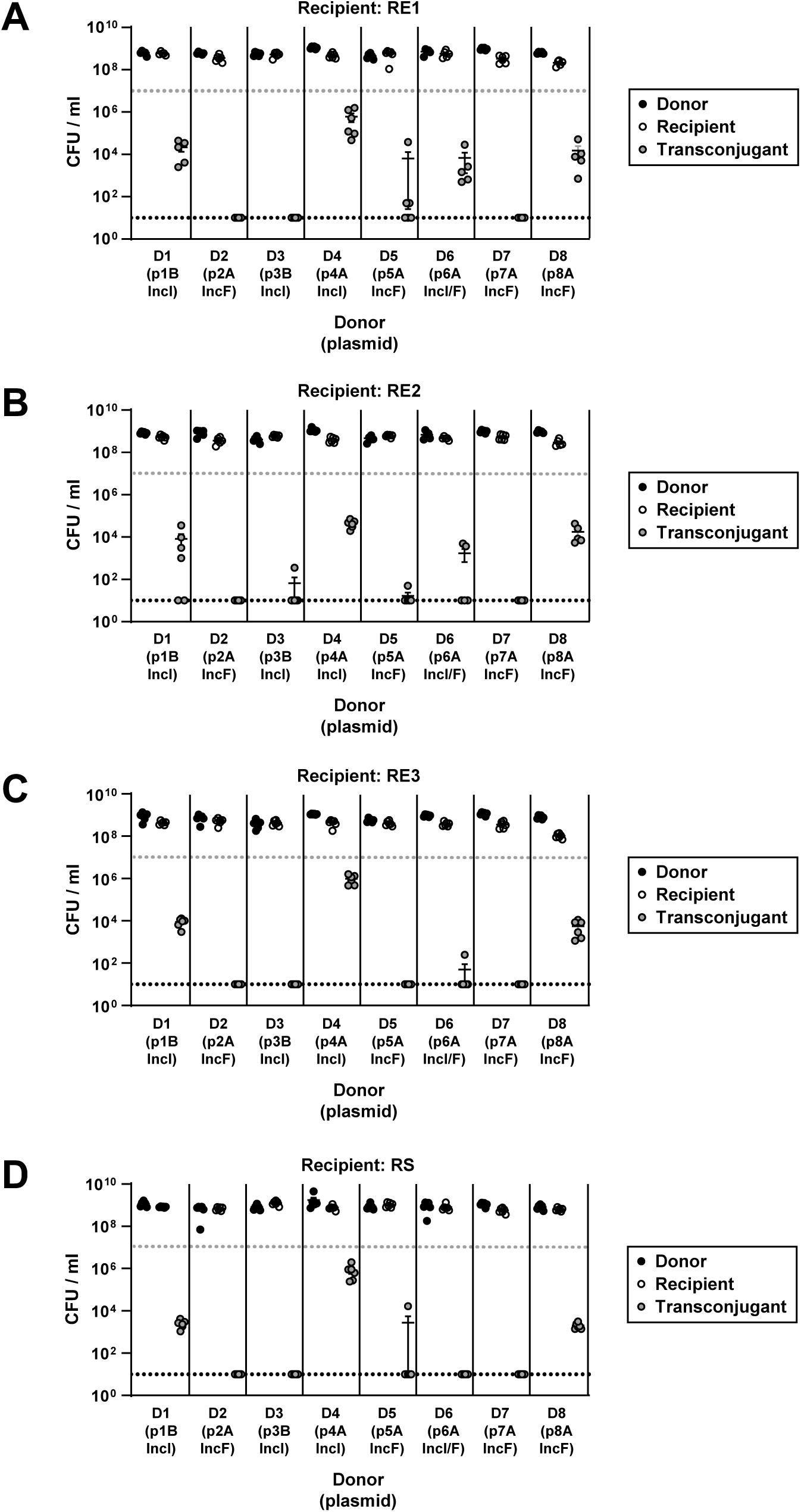
Absolute size of donor, recipient, and transconjugant populations in the 1^st^ generation *in vitro* experiment. We performed conjugation experiments with natural donor-plasmid pairs and recipients RE1 (A), RE2 (B), RE3 (C) and RS (D). Beams are mean values ± SEM (*n = 6)*, dotted lines indicate the detection limit by selective plating, grey for donors and recipients, black for transconjugants.

**Figure S11.**
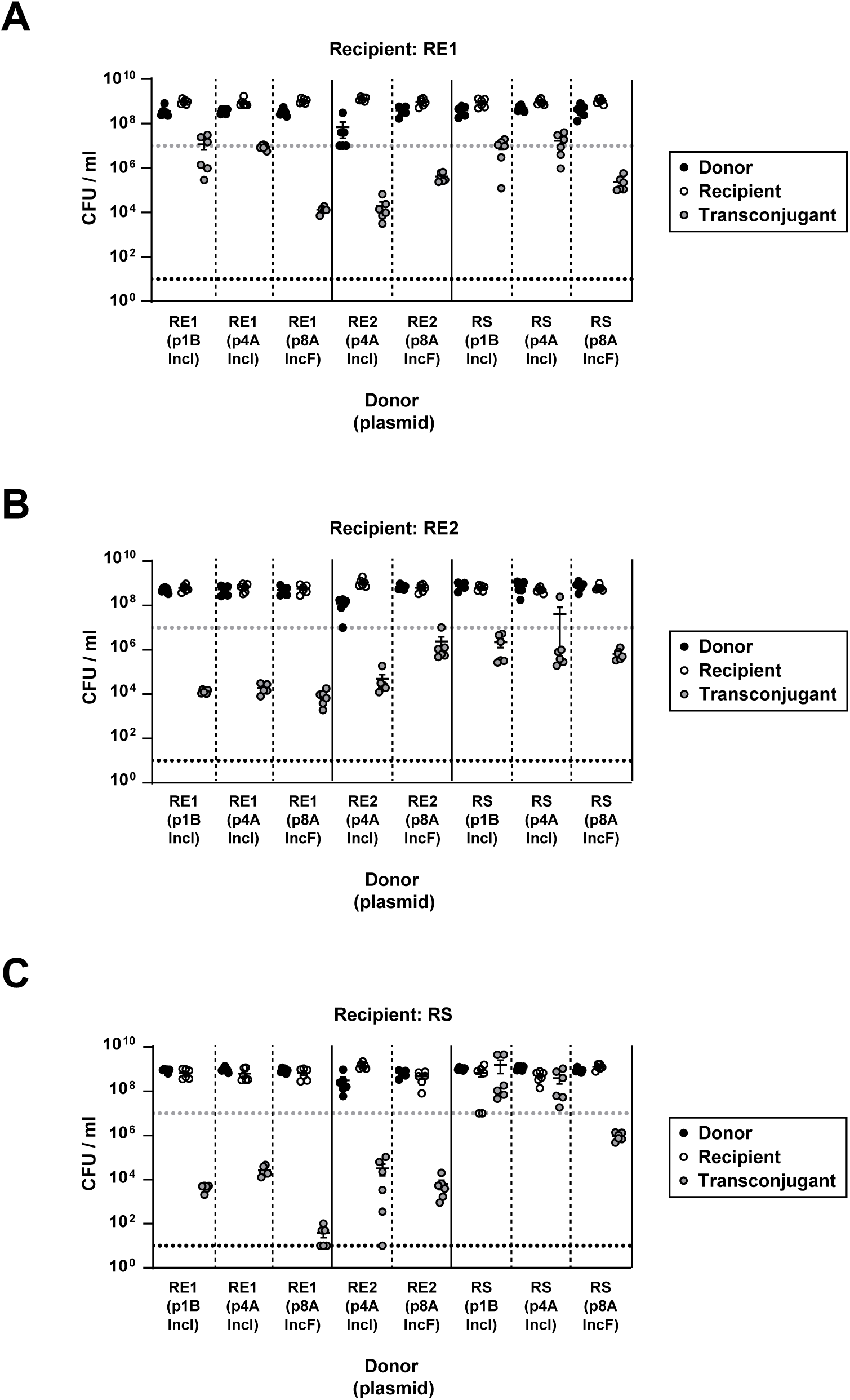
Absolute size of donor, recipient, and transconjugant populations in the 2^nd^ generation *in vitro* experiment. We performed conjugation experiments with transconjugants from the 1^st^ generation *in vitro* experiment as plasmid donors and recipients RE1 (A), RE2 (B,) and RS (C). Beams are mean values ± SEM (*n = 6*), dotted lines indicate the detection limit by selective plating, grey for donors and recipients, black for transconjugants.

**Figure S12.**
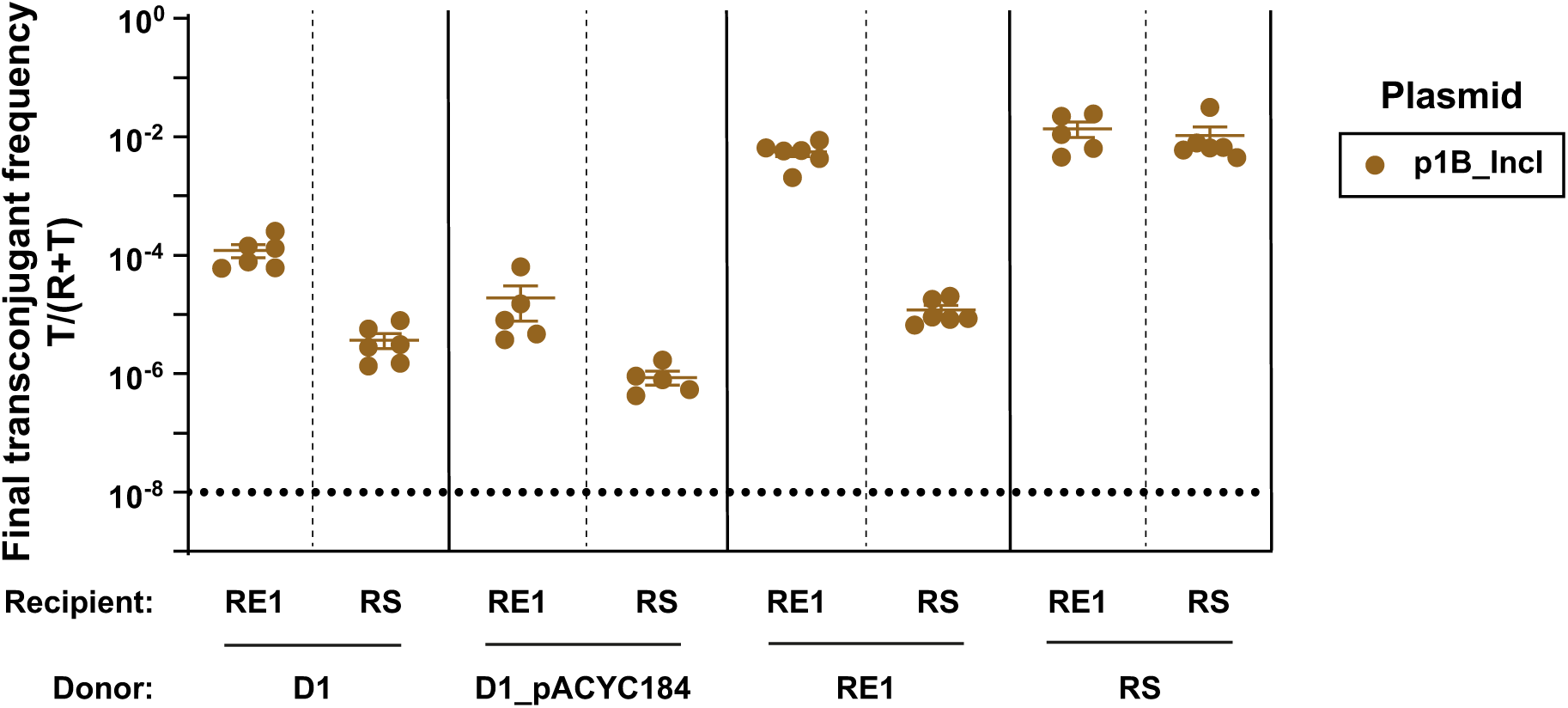
Plasmid transfer from native versus secondary host. We marked D1 with the same Cm-resistance plasmid (pACYC184) as the *E. coli* recipients in the 1^st^ generation *in vitro* experiment, to exclude pACYC184 having a major effect on transconjugants’ ability to donate plasmids. Indeed, pACYC184 in D1 had a significantly negative effect on transfer of p1B_IncI to RE1 (Wilcoxon Rank Sum Test, *P* = 0.035 after Holm’s correction for multiple testing) and to RS (Student’s t-Test, *P* = 0.042 after Holm’s correction for multiple testing). Although significant, the effect of pACYC184 was small compared to the difference in transconjugant frequency that resulted from transfer from native versus 2^nd^ generation donor strain. Circles represent independent replicates (*n = 5-6*) and the beams are mean values ± SEM. The detection limit was at ∼10^-8^.

**Figure S13.**
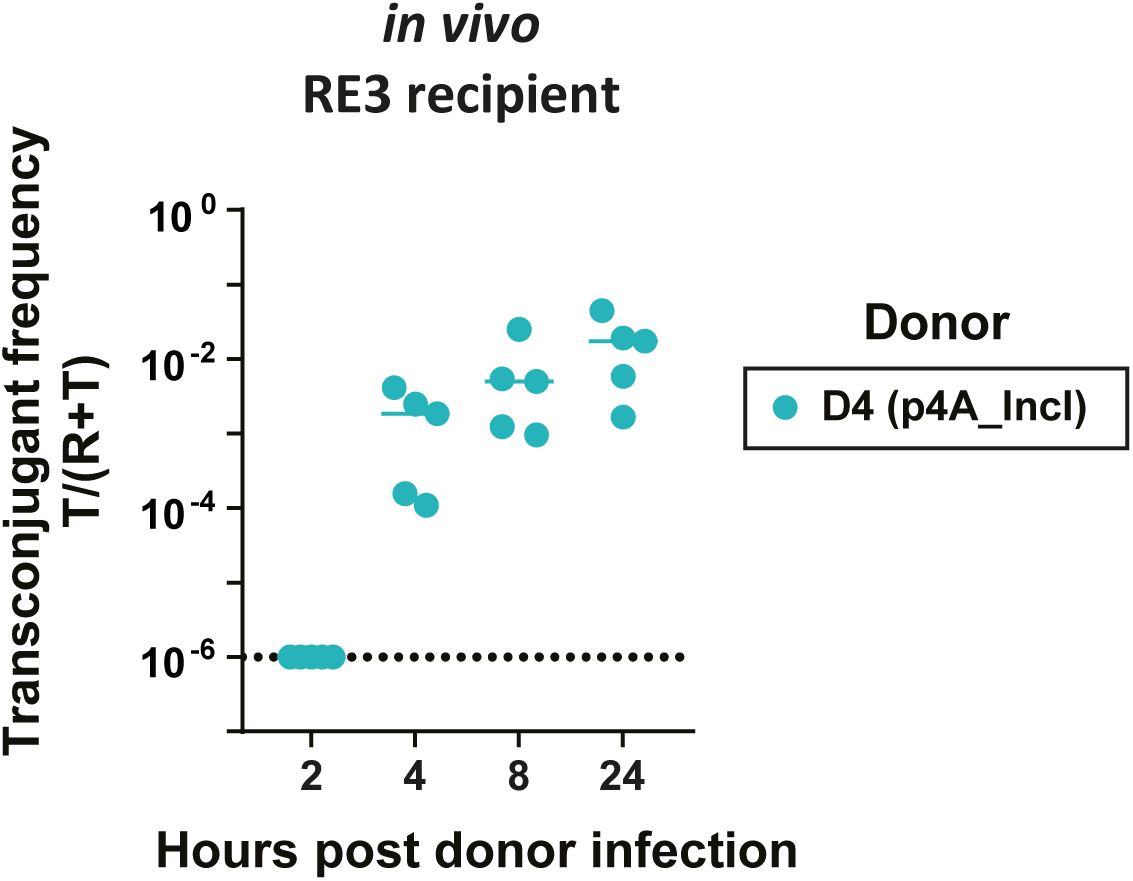
Rapid spread of p4A_IncI with recipient RE3. We monitored plasmid spread within the first 24 hours post donor infection. Plasmid spread is reported as final transconjugant frequency and enumerated in faeces by selective plating. The solid line indicates the median (*n = 5*), the dotted line indicate the detection limit for selective plating.

**Figure S14.**
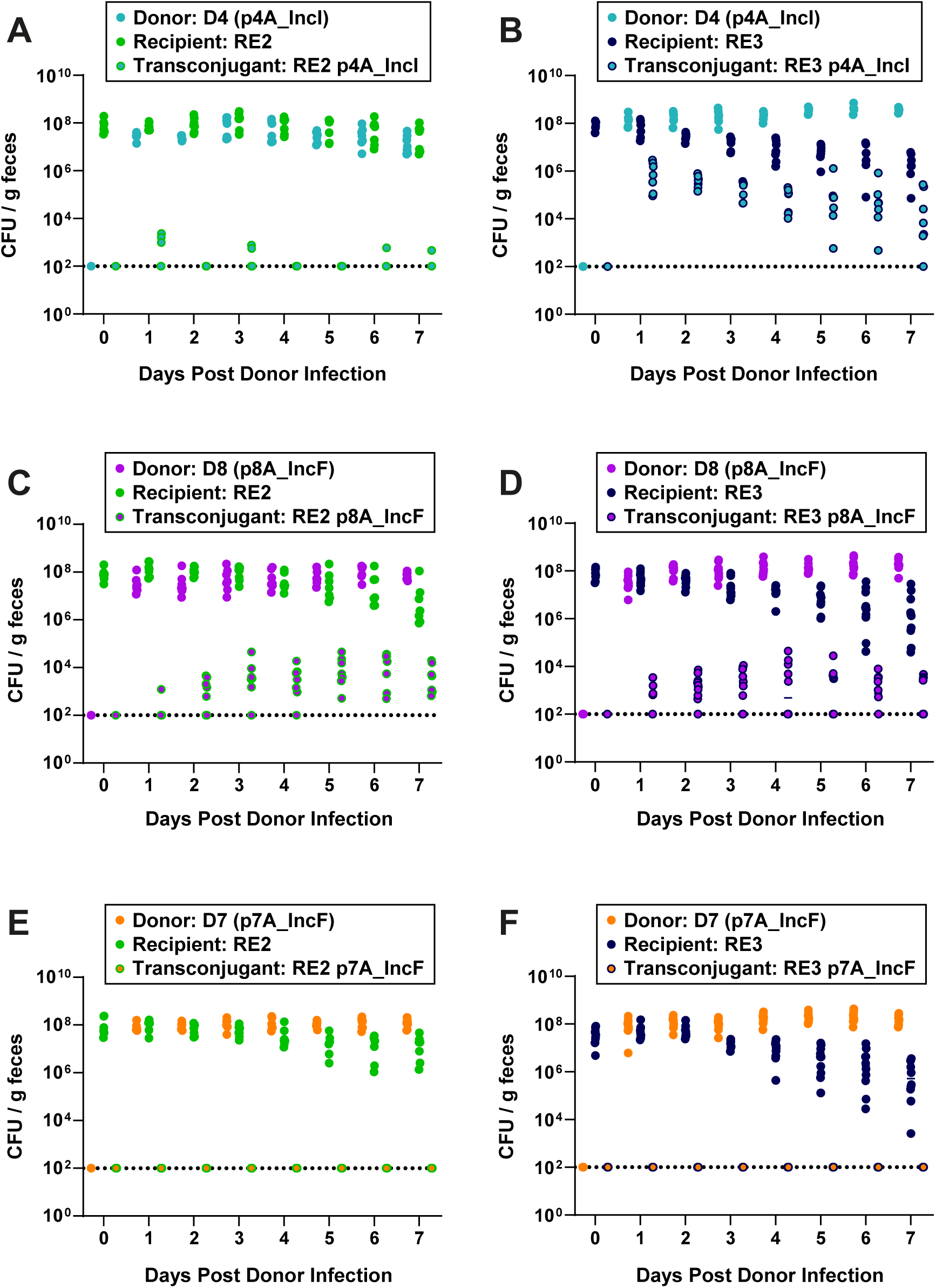
Total populations *in vivo*. Each panel reflects one donor-recipient pair and represents the population sizes of donors, recipients, and transconjugants in Fig. 4. A) Conjugation from D4 to RE2 (*n = 7*). B) Conjugation from D4 to RE3 (*n = 7*). C) Conjugation from D8 to RE2 (*n= 7*). D) Conjugation from D8 to RE3 (*n = 10*). E) Conjugation from D7 to RE2 (*n = 7*). F) Conjugation from D7 to RE3 (*n = 10*). The dotted line indicates the detection line by selective plating. The solid lines indicate the median.

**Figure S15.**
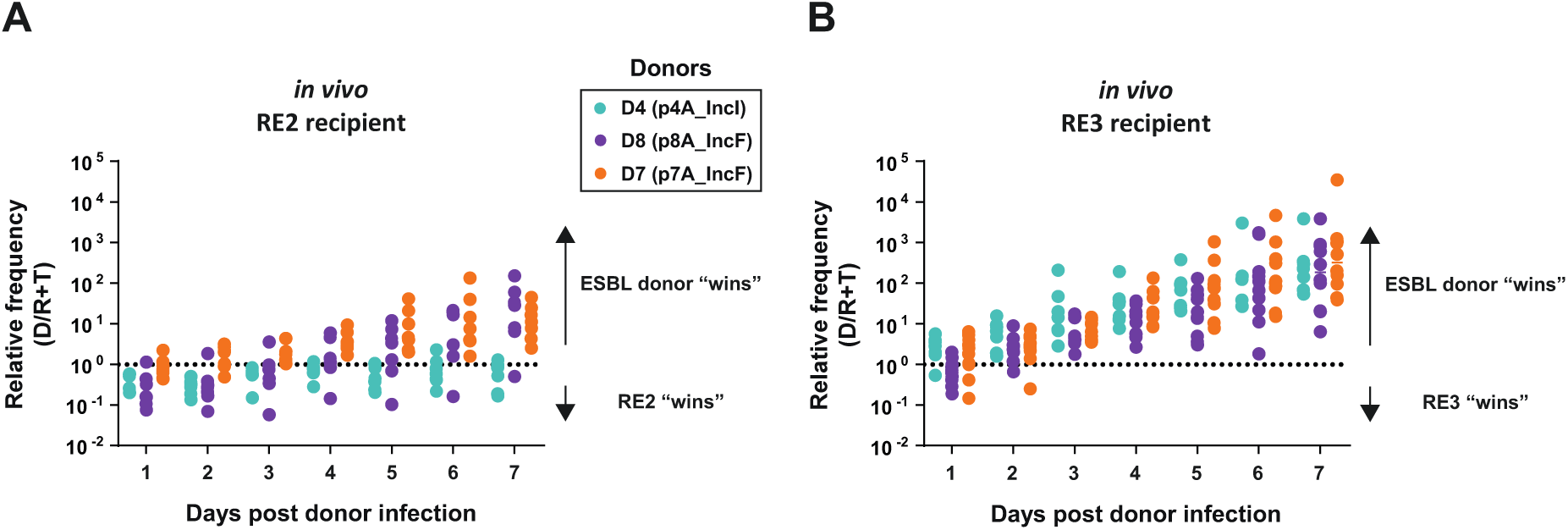
Direct competition between donors and recipients *in vivo.* We performed competition experiments by colonizing the mice with a 1:1 mix of plasmid donors with RE2 (A) (*n=7*) or RE3 (B) (*n=7* for D4-RE3; *n=10* for D8-RE3 and D7-RE3). The relative frequency was calculated by dividing the recipient population by the donor strain population.

**Figure S16.**
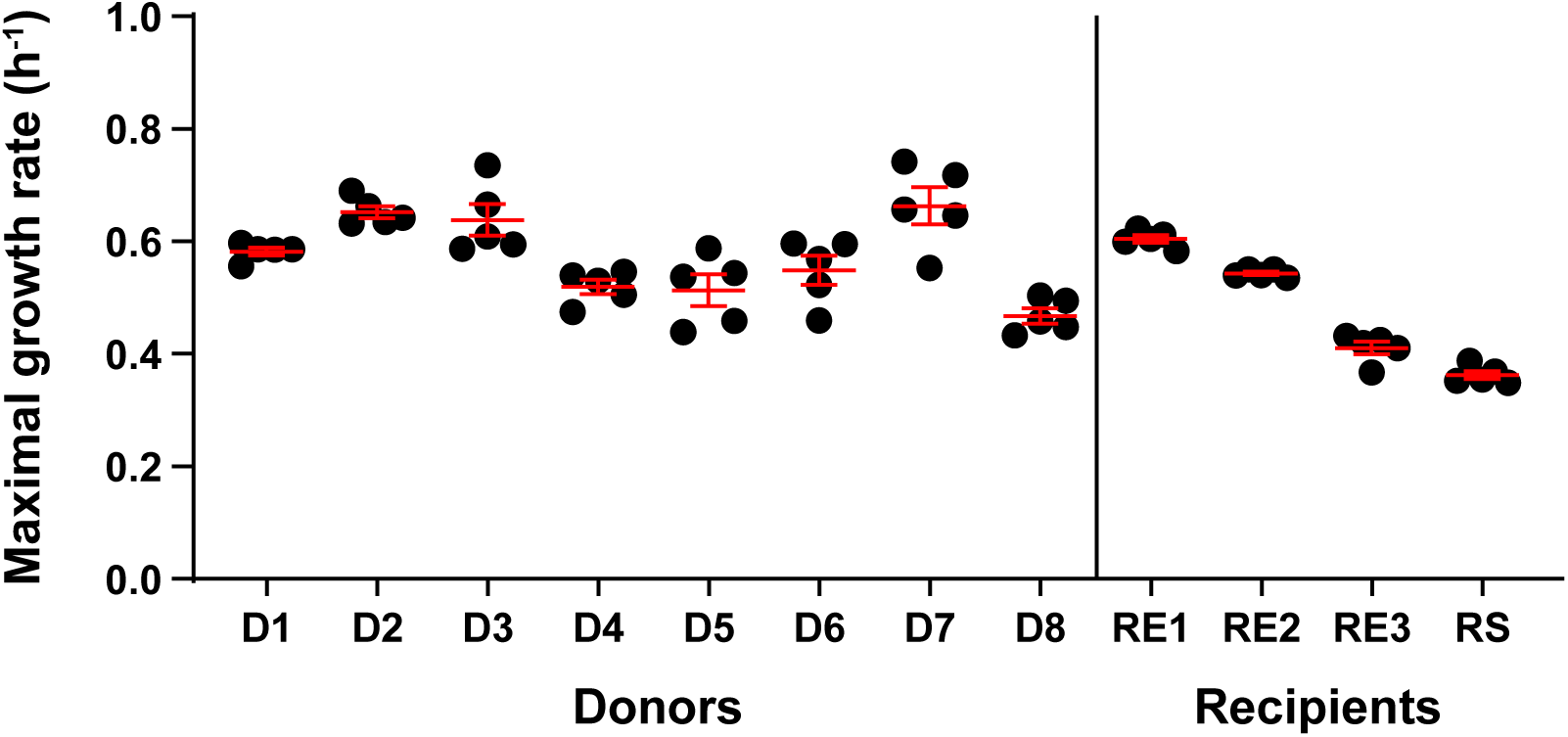
Maximal growth rates of donors and recipients used in the 1^st^ generation *in vitro* experiment. Growth rates were estimated using OD measurements over 24 hours (*n=5*) in the absence of antibiotics.

**Figure S17.**
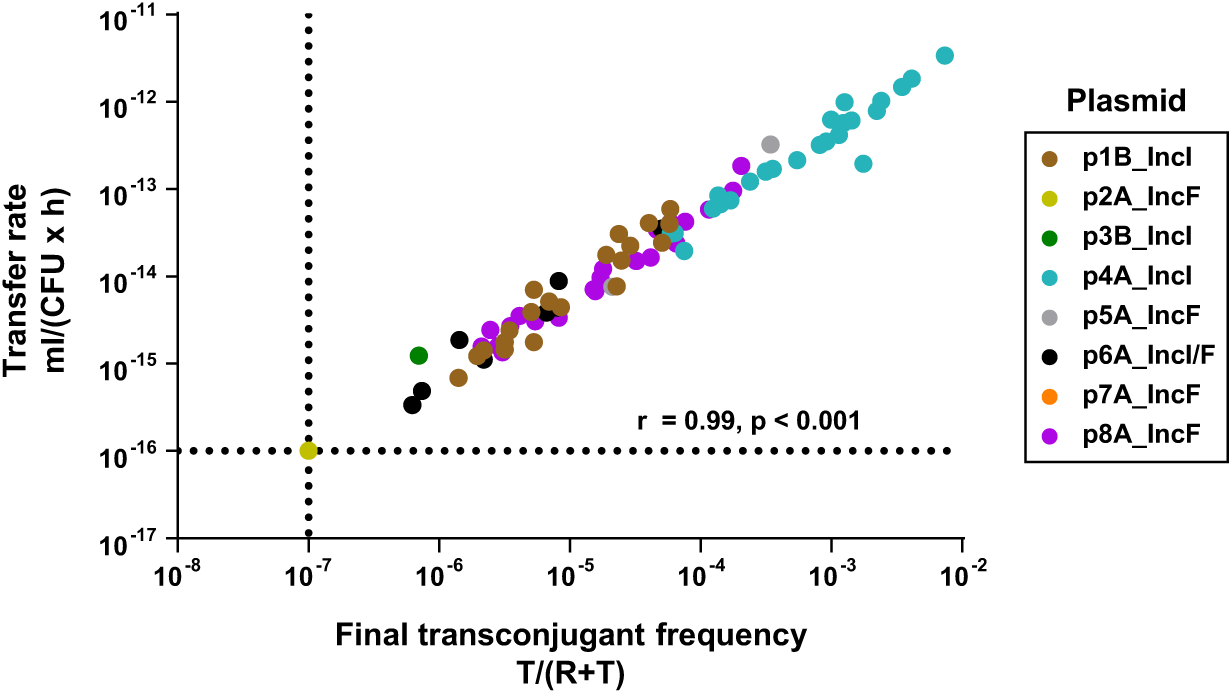
Correlation of plasmid transfer rate and final transconjugant frequency estimated using the Simonsen method. Each data point results from the same liquid mating culture shown in Fig 2. Transconjugant frequencies and transfer rates can be found in the S3 Table.

### Supplementary Tables

**Table S1 Strain overview.** Table S1 is a separate file. Overview of all strains used in this study, including their sequence type (ST), natural plasmid content and detected resistance genes. Replicon and resistance gene hits are only shown in this table if they had a coverage and percent identity of at least 70%, leading to a few differences with respect to Figures S2 and S5, most notably IncFIC_FII in D8. Contiguous sequences (contigs) were denoted “c” or “p” for chromosomal or plasmid, respectively. Short sequences (up to 5 kB) that mapped to the own chromosome, and any contigs smaller than 1kB were removed. Any remaining contigs without a known replication gene were denoted “_crypt” for cryptic.

**Table S2 Overview of plasmid transferring and mutations accumulating during conjugation experiments.** Table S2 is a separate file.

**Table S3.**
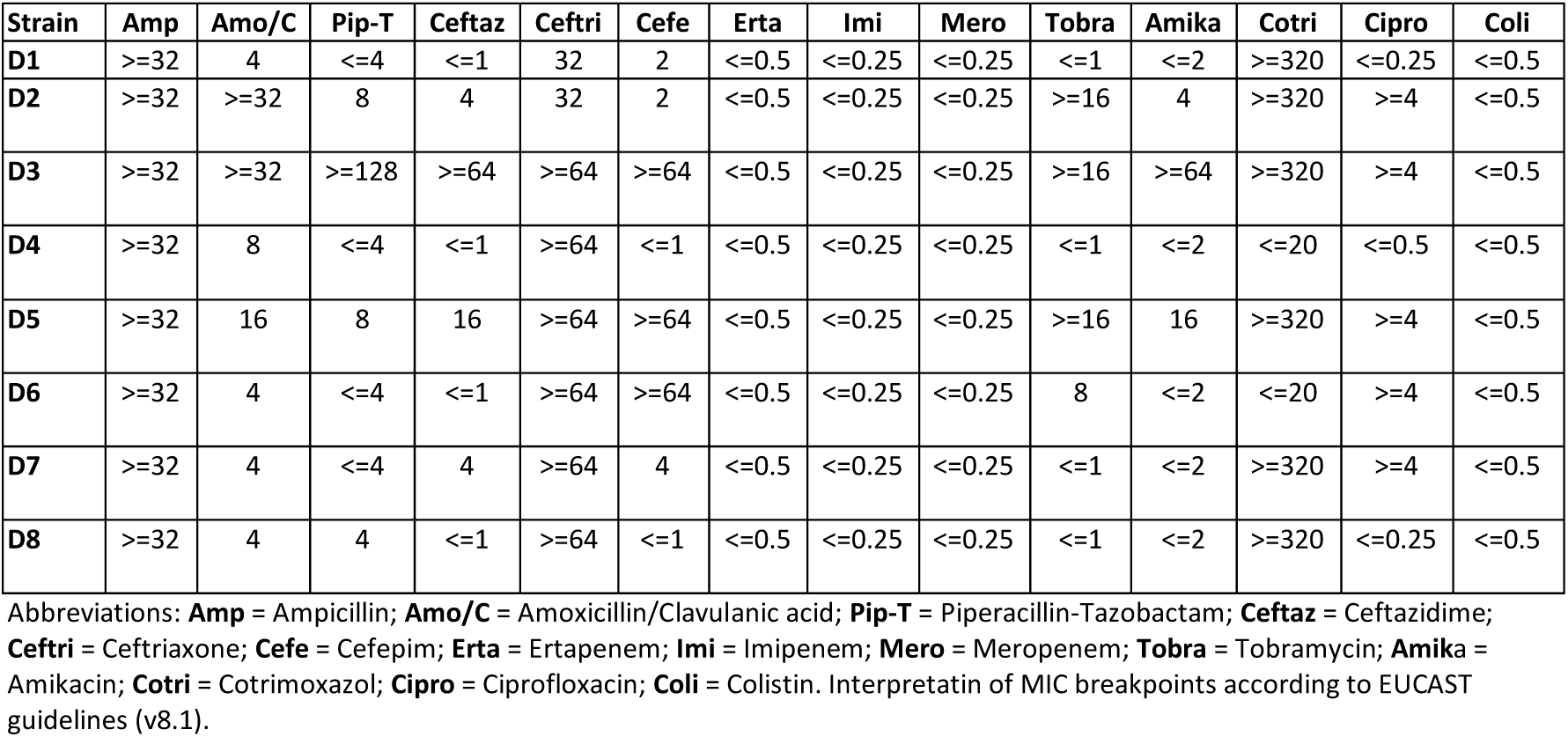
Phenotypic resistance profile of donor strains. Minimum inhibitory concentration (µg/mL) measurements of ESBL donors used in this study. The ESBL-resistance phenotype was defined by resistance to Ceftriaxone and Ceftazidime.

**Table S4.**
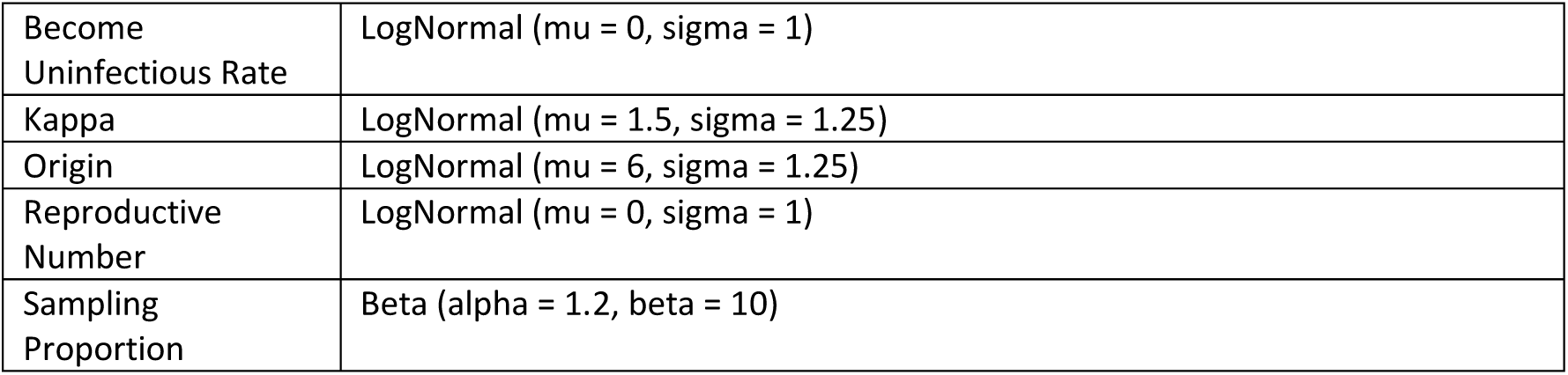
Priors for the phylogenetic tree inference. Parameter priors for the birth-death tree prior.

### Supplementary Results

#### Growth rates of all strains in the 1^st^ generation *in vitro* experiments

To calculate whether the mean difference in growth rate between recipients and transconjugants alone could explain the observed final transconjugant frequencies (Figure 2), we calculated the final transconjugant frequency that would result from such a purely clonal expansion. To simplify the calculation, we assumed exponential growth of the recipient and transconjugant population, starting from *R*(0) and a single individual *T*(0)=1 respectively. Using these assumptions, one can calculate the minimal transconjugant growth rate ψ_T_ needed to explain the final transconjugant frequency after 24 hours (*f*):

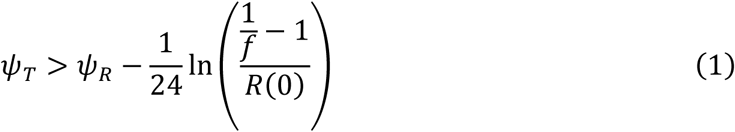

Here ψ_R_ is the corresponding plasmid-free recipient growth rate.

**Measured populations growth rates** of recipients and transconjugants. The data is the same as in the plasmid cost comparison (Figure 5A/B) and stems from manual OD measurements over 24 hours (*n = 12*). Note that these absolute growth rates differ from the ones estimates for Figures S1 and S16 (See Materials and methods).

|  | RE1_pACYC184 | RE2_pACYC184 | RE3_pACYC184 | RS_marTcat |
| --- | --- | --- | --- | --- |
| - (recipient) | $0.96 \pm 0.01$ | $0.98 \pm 0.01$ | $0.66 \pm 0.006$ | $0.34 \pm 0.002$ |
| with p1B_Incl | $0.99 \pm 0.01$ | - | - | **$0.38 \pm 0.009$** |
| with p4A_Incl | $0.95 \pm 0.01$ | $0.95 \pm 0.01$ | $0.70 \pm 0.006$ | $0.37 \pm 0.001$ |
| with p8A_IncF | $0.88 \pm 0.004$ | $0.99 \pm 0.006$ | $0.67 \pm 0.004$ | **$0.38 \pm 0.0003$** |

**Calculated minimal transconjugant growth rates** needed to explain the observed final transconjugant frequencies (Figure 2).

|  | RE1_pACYC184 | RE2_pACYC184 | RE3_pACYC184 | RS_marTcat |
| --- | --- | --- | --- | --- |
| with p1B_Incl | 1.07 | 1.06 | 0.75 | **0.36** |
| with p4A_Incl | 1.23 | 1.16 | 0.96 | 0.60 |
| with p8A_IncF | 1.10 | 1.13 | 0.80 | **0.36** |

In two cases, i.e. transconjugant RS carrying p1B_IncI and p8A_IncF, the measured growth rate of transconjugants was greater than the calculated minimal growth rate needed to explain the observed final transconjugant frequencies. Thus, for these two out of the 10 donor-recipient combinations (excluding RE2 and RE3 carrying p1B_IncI), we cannot exclude that final transconjugant frequencies were reached by clonal expansion, rather than conjugative transfer. Our calculations, however, are conservative, because of the assumption of exponential growth, which overestimates the number of transconjugants that could have resulted from clonal expansion.

